# Membrane-Dependent Dynamics and Dual Translocation Mechanisms of ABCB4: Insights from Molecular Dynamics Simulations

**DOI:** 10.1101/2024.10.20.618436

**Authors:** Veronica Crespi, Ágota Tóth, Angelika Janaszkiewicz, Thomas Falguières, Florent Di Meo

## Abstract

ABCB4 is an ATP-binding cassette transporter expressed at the canalicular membrane of hepatocytes and responsible for translocating phosphatidylcholine into bile. Despite the recent cryo-EM structures of ABCB4, knowledge about the molecular mechanism of phosphatidylcholine transport remains fragmented. In this study, we used all-atom molecular dynamics simulations to investigate ABCB4 dynamics during its transport cycle, leveraging both symmetric and asymmetric membrane models. Our results demonstrate that membrane composition influences the local conformational dynamics of ABCB4, revealing distinct lipid-binding patterns across different conformers, particularly for cholesterol. We explored the two potential mechanisms for phosphatidylcholine translocation: the canonical ATP-driven alternating access model and the “credit-card swipe” model. Critical residues were identified for phosphatidylcholine binding and transport pathway modulation, supporting the canonical mechanism while also indicating a possible additional pathway. The conformer-specific roles of kinking in transmembrane helices (TMH4 and TMH10) were highlighted as key events in substrate translocation. Overall, ABCB4 may utilize a cooperative transport mechanism, integrating elements of both models to facilitate efficient phosphatidylcholine motion across the membrane. This study provides new insights into the relationship between membrane environment and ABCB4 function, contributing to our understanding of its role in bile physiology and susceptibility to genetic and xenobiotic influences.

## 1. Introduction

ABCB4 is an ATP-binding cassette (ABC) transporter exclusively expressed at the canalicular membrane of hepatocytes. It is responsible for the translocation of phosphatidylcholine lipids (PC) into the bile canaliculus; PC being one of the main components of bile’s mixed micelles, along with bile salts and cholesterol^1^. Variations in *ABCB4* gene are associated with biliary diseases such as progressive familial intrahepatic cholestasis type 3 (PFIC3), cholelithiasis syndrome, intrahepatic cholestasis of pregnancy, and liver cancer^2^. Xenobiotics can inhibit ABCB4 leading to iatrogenic imbalance of bile composition which is associated with an increment in bile toxicity owing to bile salt detergent action on cell membranes ^3–5^.

Recently, ABCB4 structure was resolved through cryogenic electron microscopy (cryo-EM) adopting the so-called type IV fold^1,3^. ABCB4 structure assembles two consecutive similar subdomains connected by a linker ^6^. Each subdomain consists of a transmembrane domain (TMD) made of six transmembrane helices (TMHs) and one so-called nucleotide binding domain (NBD). ABCB4 was resolved under different conformations, namely inward facing (^IF^ABCB4) ^3^ and close-cleft (^CC^ABCB4) ^1^. Different bound states were resolved, namely apo, substrate- and inhibitor-bound states for ^IF^ABCB4 ^3^, and nucleotide-bound state for ^CC^ABCB4^1^. Following the commonly accepted two-side alternating access model, both the NBD dimerize upon the binding of two ATP molecules sandwiched in two nucleotide binding sites (NBS) ^7^. It is still not fully understood how a PC substrate is translocated during the transport cycle. Likewise, knowledge remains fragmented regarding whether ABCB4-mediated PC lipid is directly released to canalicular micelles or if ABCB4 transport mechanism leads to outer leaflet PC enrichment prior to PC lipid extrusion in bile, even though the latter seems more likely^8^.

Two complementary different molecular mechanisms have been suggested regarding PC translocation across lipid bilayer. Nosol et al. proposed the canonical ATP-driven alternating access for PC translocation relying on the resolution of substrate-bound ABCB4 adopting the IF conformation^3^. Similar to type IV ABC-driven substrate translocation^3^, ^IF^ABCB4 might initially bind PC lipid substrate from the inner leaflet leading to the ligand-bound conformation (^IF^ABCB4-PC). PC substrate is thus expected to enter into ABCB4 binding pocket either by the front gate composed of TMH3/TMH4/TMH6 or the rear (or back) gate formed by TMH9/10/12, the latter being suggested to be less likely^9^. The central role of Trp234 and Phe345 for PC substrate binding was proposed through site-directed mutagenesis experiments and the cryo-EM structure. Upon ATP binding, ^IF^ABCB4-PC-(ATP)_2_ state might undergo large-scale conformational changes leading to PC release out of the protein binding pocket through a proposed transient open outward-facing (OF) conformation.

Interestingly, site-directed mutagenesis of Trp234, Ala231, Phe345, or His989 was associated with a significant decrease of PC translocation ranging from ca. 20 to 50% as compared to WT-ABCB4; but no total inhibition of ABCB4 function was observed^3^. This may suggest the existence of a non-canonical PC translocation mechanism. In this context, a joint computational and experimental study proposed an alternative ABCB4-driven PC translocation adopting the so-called credit-card (c/c) swipe mechanism^9^. Such a mechanism relies on the exposure of TMH1 polar amino acids shading PC polar head while maintaining lipid tails in a hydrophobic environment. The c/c swipe mechanism starts with the outward-open or ^CC^ABCB4 conformation, in which TMH1 polar amino acids bind PC polar head. As for other type IV ABC transporters^6,10^ ATP hydrolysis is required for CC-to-IF conformational transition, achieving PC translocation. Subsequently, the binding of ATP in NBS favors conformational changes from ^IF^ABCB4-(ATP)_2_ back to ^CC^ABCB4-(ATP)_2_.

Taking advantage of the available cryo-EM structures of apo ^IF^ABCB4, ^IF^ABCB4-PC and ^cc^ABCB4, we investigated the dynamics of the main conformational milestones by all-atom molecular dynamics (MD) simulations. Nucleotide-bound state was also considered for ^IF^ABCB4 conformations, namely ^IF^ABCB4-(ATP)_2_ and ^IF^ABCB4-PC-(ATP)_2_. In parallel, lipid environment has been shown not only to act as stabilizing physical containers of membrane proteins but also to probably play an active role in *e.g.*, G-protein coupled receptor^11^, major facilitator superfamily proteins^12^ and ABC transporters^13–18^. In other words, lipid membrane composition is thus of particular importance for the modulation of membrane protein functions. Over the past decades, MD simulations were often carried out in simplistic and symmetric membrane models mostly consisting of 1-palmitoyl-2-oleyl-sn-glycero-3-phosphatidylcholine (POPC), 1-palmitoyl-2-oleyl-sn-glycero-3-phosphatidylethanolamine (POPE) and/or cholesterol (Chol). Nevertheless, it is now possible to model more realistic membranes thanks to the tremendous recent advances made in lipid force fields (*e.g.*, CHARMM, SLipids, Lipid21^19–21^). In this context, special attention is thus also paid to the impact of the lipid composition onto the structural dynamics of ABCB4. Two different membrane models were employed, namely symmetric (POPC:POPE:Chol (2:1:1)) and asymmetric distributions consisting in multiple lipids for which proportions were inspired from Ingólfsson et al.^22^. Lipid model compositions are reported in Table S1 and S2. In this asymmetric model, we also include phosphatidylserine (PS) and phosphatidic acid (PA) polar head lipids. We also include other unsaturated lipid tails such as arachidonate and docosahexaenoate, leading to a more realistic membrane model (See Table S2). The present study aims to provide computational perspectives on membrane-dependent ABCB4 structural dynamics as well as the complementarity between the canonical substrate translocation and the credit-card swipe mechanism.

## 2. Materials & Methods

### 2.1. Structure

ABCB4 five main states and conformations along canonical transport cycle were considered, namely: apo ^IF^ABCB4, ^IF^ABCB4-PC, ^IF^ABCB4-(ATP)_2_, ^IF^ABCB4-PC-(ATP)_2_, and ^CC^ABCB4-(ATP)_2_. ^IF^ABCB4-based models were built using either apo or PC-bound cryo-EM structures (PDB ID: 7NIU, 7NIV) while ^CC^ABCB4 was constructed using the resolved cryo-EM structure of ABCB4 adopting CC conformation (PDB ID: 6S7P). The so-called linker 1 connecting the NBD1 to TMH7 was not included in the present molecular model given its expected disordered structure to preclude false non-covalent interactions with resolved domains. Few residues of the first extracellular loop connecting TMH1 and TMH2 being unresolved, missing residues (*i.e.*, from Phe92 to Gly106 for ^IF^ABCB4 structures - *i.e.*, 7NIU and 7NIV – and Lys84 to Pro105 ^IF^ABCB4 – *i.e.*, 6S7P) were included using the Modeller v9.23 ^23^ taking advantage of resolved P-gp structures (PDB ID: 6QEX) ^18^. The cryo-EM resolved 1,2-dilinoleoyl-sn-glycero-3-phosphatidylcholine (DLPC) lipid was used for substrate-bound states of ^IF^ABCB4 conformations (*i.e.*, ^IF^ABCB4-PC and ^IF^ABCB4-PC-(ATP)_2_. Since resolved ^IF^ABCB4 structures do not include ATP and Mg^2+^ molecules, ATP-bound states were built by independently superimposing ^IF^ABCB4 NBDs onto those obtained from ^CC^ABCB4 conformation in which ATP and Mg^2+^ were expected to be properly docked to at least NBD A-loop and signature motifs. Protonation states of titratable amino acids were predicted using the propKa3.0 software^24,25^ Histidine residues were protonated by visual inspection of their potential involvement in H-bond network with surrounding amino acids. For sake of comparison, the same protonation states were used for each model.

Molecular models were briefly minimized in vacuum to remove potential steric clashes using the Amber 20 package^26,27^ prior to embedding into lipid bilayer membranes using the CHARMM-GUI web server^28–30^. Models were first carefully aligned with ^IF^ABCB4 and ^CC^ABCB4 coordinates in the OPM (Orientations of proteins in membranes) database^31^ to mimic proper protein alignment in membrane models. Two models of lipid bilayers were here considered to investigate the protein-lipid interplay and the dependence on membrane composition. A classical but simplistic symmetric lipid bilayer made of 1-palmitoyl-2-oleyl-sn-glycero-3-phosphatidylcholine (POPC), 1-palmitoyl-2-oleyl-sn-glycero-3-phosphatidylethanolamine (POPE) and cholesterol (Chol) with a molecular ratio of POPC:POPE:Chol (2:1:1). Besides, a more complex asymmetric lipid bilayer composition was also considered made of different lipids. PC, PE, Chol but also phosphatidic acid (PA) and phosphatidylserine (PS); lipid polar heads were built as well as different lipid tails such palmitoyl (PA, 16:0), oleoyl (OL, 18:1), arachidonyl (AR, 20:4), docosahexaenoyl (DHA, 22:6). Lipid compositions in inner and outer leaflet for each model are reported in Table S1 and S2. Overall, the number of lipids in all the membranes is slightly larger than 400 leading to system size of ca. 120x120x180 Å^3^ including water and NaCl counterions set up at the extracellular physiological concentration (*i.e.*, 0.15 M, system details are reported in Table S3). The scripts included in AmberTools package (namely, charmmlipid2amber.py and pdb4amber) were employed to convert the CHARMM-GUI inputs in an Amber-compatible file format. For bound states, PC substrate, ATP molecules, and Mg^2+^ ions were then added by superimposing models built initially without lipid bilayer. For each system, the corresponding amount of counterions was removed to maintain the neutrality of the system.

### 2.2. Molecular dynamics simulations

Amino acids, lipids, water, and ATP molecules were respectively modelled using the FF14SB^32^, lipid21^21^, TIP3P^33–35^, and DNA.OL15-based forcefields^36^ for bound models. Apo ^IF^ABCB4 systems were modelled using the latest FF19SB^37^ with OPC^38^ model for water molecules. Such a difference of forcefield models between apo and bound states was due to unexpected forcefield incompatibilities. Using FF19SB and OPC forcefield with substrates surprisingly leads to unreliable restraints onto amino acid backbone and sidechains. Mg^2+^ ions and NaCl counterions were modeled using the mono and divalent parameters suited either to TIP3P^39^ or OPC ^38^ water force fields, with FF14SB or FF19SB, respetively. ABCB4 PC substrate (*i.e.*, DLPC) was modelled using the lipid21 forcefield. The cutoff for non-bonded interaction was set at 10 Å for both Coulomb and van der Waals potentials. Long-range electrostatic interactions were computed using the particle mesh Ewald method ^40^.

Simulations were conducted using the Amber20 and Amber22 packages^41,42^ using CPU and GPU versions of the PMEMD code. First, minimizations using energy steepest gradient were conducted as follows: (i) minimization of water O-atoms for 20,000 steps, (ii) all bonds involving H-atoms for 20,000 steps, (iii) water molecules and counterions for 50,000 steps, and (iv) the whole system for 50,000 steps. For each model, system thermalizations were performed using Langevin thermostat in two steps. First, water molecules were thermalized from 0 to 100 K for 50 ps in the NVT ensemble with 0.5 fs as time integration. Then, the whole system was heated up from 100 K to 310 K for 500 ps in the NPT ensemble using a time integration of 2 fs, and restraining bonds involving H-atoms with the SHAKE algorithm. Pressure and box equilibrations were then achieved by performing short MD simulations for 5ns with a integration time set at 2 fs in the semi-isotropic NPT ensemble for which Berendsen barostat ^43^ was employed. Finally, for each model, MD productions of 2 µs were performed using 2 fs as integration time step, in the NPT ensemble condition with semi-isotropic scaling. Langevin dynamics thermostat ^44^ with 1.0 ps^-1^ collision frequency was used to maintain the temperature. The overall pressure was maintained at 1 bar with semi-isotropic pressure scaling using Berendsen barostat.

Noteworthy, ATP-bound models were carried out using a restraint MD approach as used for ABCC1 transporter ^13^ and initially proposed by Wen et al. in 2013 ^45^ to ensure the different induced fit for ATP-Mg^2+^-NBD interactions. Briefly, for equilibration steps, distance between each ATP purine moiety and NBD1&2 A-loop tyrosines (namely Tyr 403 and Tyr 1043) were restrained as well as H-bond between phosphate moieties, Mg^2+^ atoms with surrounding Walker A serine residues (Ser436 and Ser1076) and Q-loop glutamine residues (Gln477 and Gln1206) to maintain proper ATP docking in NBS. For each restraint, a harmonic potential was applied to restrain distances using ATP binding modes of ^cc^ABCB4-(ATP)_2_ as reference. The distance restraints were applied during the thermalization, equilibration, but also the first 10 ns of the production. They were then smoothly removed over 10 ns. The remaining part of the production run was performed without restraints. Distances were then monitored and showed in Supplementary Figures S1-S4. Even though distance may differ from initial restraints, ATP molecules remain in NBSs along MD simulations. For each model, three replicas were conducted from the thermalization stage to consider a minimum of conformational variability. The aggregated total MD simulation time in the present work is ∼60 µs (see Table S4).

### 2.3. Analysis and visualization

For each system and given time-dependent root-mean square deviations (RMSD) evolution over MD simulations (see Figure S5), analyses were performed over the last 1 μs of each production run. The maximum observed for the replica 1 of ^IF^ABCB4-PC-(ATP)_2_ was due to a transient NBD dimerization at the beginning of the trajectory during the equilibration prior to NBD reopening after 1 μs. Trajectories were saved every 100 ps. Analyses were conducted with the CPPTRAJ^46^ and MDAnalysis^47,48^ softwares as well as in-house python scripts. Plots were created using the matplotlib v3.1.1 python package ^49^. Trajectories visualization and image rendering were made using the VMD software (versions 1.9.3 and alpha-v1.9.4)^50^.

The so-called ABC structural parameters, namely, intracellular (IC) and extracellular (EC) angles, NBD distance, and NBD twist were used to assess the conformational difference of the different models. They were defined according to previous studies by Moradi et al.^51^, Hofmann et al.^52^, and Tóth et al.^13^ (see Figure 1A). Briefly, IC was defined as the angle between vectors made of the EC TMH region center-of-mass towards the centers-of-mass of IC TMH regions bound to either NBD1 (*i.e.*, TMH1, TMH2, TMH3, TMH10, TMH11, and TMH6) or NBD2 (*i.e.*, TMH4, TMH5, TMH7, TMH8, TMH9, and TMH12). Likewise, EC angle was defined by the angles originating from the center-of-mass of intracellular regions of TMHs towards the extracellular regions of either TMH1, TMH2, TMH11, TMH12 or TMH5, TMH6, TMH7. The NBD distance was defined by the distance between the centers-of-mass of each NBD; and finally, the NBD twist measures the dihedral angle between the two NBDs, defined by four centers-of-mass made of alpha-helical subdomains and beta-sheet subdomain of each NBD. The TMH and NBD topology used for ABCB4 protein is reported in Table S5. Residues selected for ABC structural parameter calculations are reported in Supporting Table S6. Free energy surfaces were estimated using the InfleCS method (Inflection core state Clustering)^53^. Briefly, using user-defined structural parameters, distributions are fitted to Gaussian mixture model-based density to identify local minima and therefore potential metastable core states. This was conducted using either ABC structural parameters (i.e., IC angle, NBD distance and NBD twist) to assess the protein conformational dynamics or PC polar head orientation and PC distance to membrane center to investigated substrate dynamics in ABCB4 binding pocket. Structural principal component analyses (PCA) were also performed with the CPPTRAJ package^46^, either focusing on the ABC core (*i.e.*, backbone atoms of TMH1 to TMH12 and NBD1 and NBD2) or only TMHs. In the first case, the system variability was measured by considering each replica per system independently aligned to an average structure of the system. In the second case, PCA was carried out by aligning all the systems to an average structure of the ^CC^ABCB4 conformation to measure the overall variability along the transport cycle. For PC-bound states, contacts between PC substrate and ABCB4 were investigated using the CPPTRAJ package. Lipid order was calculated with the CPPTRAJ package using the keyword *lipidscd*. Polar-head oriented lipid distributions and residencies^54,55^ were measured using the CPPTRAJ package (*grid* keyword with a resolution of 1 Å). For each polar head or entire cholesterol, the threshold of lipid occupancies was set up to highlight regions for which the likelihood of finding a given polar head was higher than 60% over the analysis. Polar head and cholesterol contacts have been investigated by CPPTRAJ package. Residency time was calculated using the ProLint software^54^ (version 2.0, https://github.com/ProLint/prolint2) from contact analyses using a 7 Å cutoff distance.

**Figure 1.**
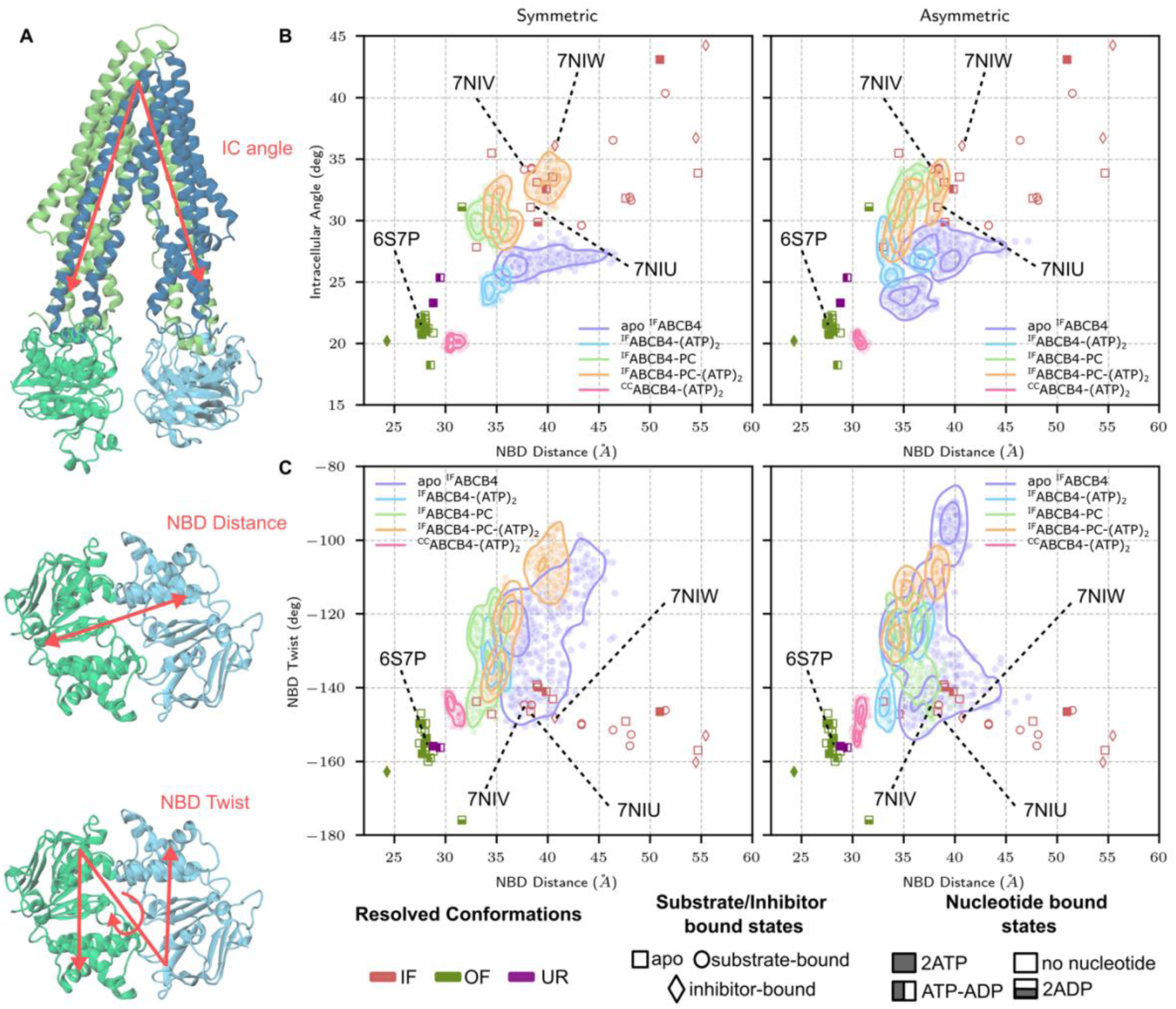
Projection of ABCB4 MD simulations onto the ABC conformational space. (A) Definitions of the ABC structural parameters, TMH1-6 and NBD1 are depicted in green and TMH7-12 and NBD2 are depicted in blue. Conformation- and bound state-dependent distributions of (B) IC angle *versus* NBD distance and (C) NBD twist *versus* NBD distance from ABCB4 MD simulations on the ABC conformational space defined by resolved cryo-EM structures in symmetric and asymmetric models.

## 3. Results

### 3.1. Membrane-dependence of ABCB4 structural dynamics

#### 3.1.1. Overview of the ABC conformational space along transport cycle

Sampling of the ABCB4 structural dynamics was examined accounting for the different conformations (namely IF and CC) and bound states (PC-, ATP- and PC/ATP-bound). The so-called ABC structural parameters^51,52^ *i.e.*, intracellular (IC) angle, NBD distance, and NBD twist were monitored throughout the MD simulations (Figure 1A). These structural parameters were used to investigate structural variabilities of IC opening as well as NBD dimerization and twist along ABCB4 alternating access^51^. Distributions of ABC structural parameters were projected onto the so-called ABC conformational space (see Figure 1B&C). It was defined by ABC structural parameters measured from a large dataset of cryo-EM structures of type IV ABC transporters (including the ABCB4 structures) and adopting different conformations and bound states (Table S7). The three main conformations (namely IF, OF, and the so-called Unlocked-returned – UR) are neatly arranged and recognizable, even though IF structures exhibit large variability regarding IC angle and NBD distance. Ligand- and/or nucleotide-bound states are less clearly distinguishable.

Interestingly, for most of the IF ABC conformations, substrate-bound structures reported larger IC angles than apo states in line with our MD simulations (Figure 1B). Inhibitor-bound ABC IF conformations exhibited an even larger IC opening. This is the case for ^IF^ABCB4 structures (PDB IDs: 7NIU, 7NIV, and 7NIW being respectively apo, substrate- and inhibitor-bound states, see Figure 1B). While comparing structures of ABCB4 and those obtained from MD simulations, similar behaviors were observed regardless of membrane composition. ABC structural parameters measured from ABCB4 structures exhibit only slight differences with MD simulations, in contrast to similar investigations conducted with the multidrug resistance-associated protein 1 (ABCC1/MRP1)^13^. Cryo-EM IF structures of ABCB4 exhibit slightly larger IF opening as pictured by IC angle and NBD distances as compared to MD simulations. In contrast, structures relaxed during MD simulations, leading to decrease in NBD distance and IC angle for IF conformations. NBD distance slightly increased during the MD simulations for ^CC^ABCB4-(ATP)_2_ while NBD twist remained within the same range as cryo-EM structure. As expected, owing to the absence of substrate or nucleotide, apo states exhibit much larger structural variability of IC opening as pictured by NBD distance and IC angle distributions, in agreement with observations made on *e.g.*, MRP1^13^ or P-gp^10^.

Regardless of membrane composition, MD simulations conducted with IF conformations show that ATP binding is not sufficient to reach the so-called proper NBD dimerization described by the ^CC^ABCB4-(ATP)_2_ conformation (Figure 1B&C). This can be explained by the relatively short μs-scale from all-atom MD simulations as compared to the expected alternating access timescale, in line with observations made on other ABC transporters ^10,13^. This also suggests that IF-to-OF transition requires slow large-scale motions driven from TMHs to trigger proper NBD dimerization. Indeed, IF conformations showed NBD distance values greater than those of the CC conformations (30.8 ± 0.4 and 30.7 ± 0.3 Å for ^CC^ABCB4-(ATP)_2_ respectively in symmetric and asymmetric membrane). Likewise, MD simulations revealed significantly lower variability of NBD twist for ^CC^ABCB4-(ATP)_2_ conformation (-145.4° ± 2.0 and -149.1° ± 3.3 in symmetric and asymmetric membrane, respectively).

Binding of PC substrate in the canonical binding pocket is associated with an increase in the IC opening as compared with apo ^IF^ABCB4 state likely owing to the steric hindrance assigned to PC substrate size. In contrast, NBD distances for ^IF^ABCB4-PC tend to be smaller than apo ^IF^ABCB4 state, suggesting an allosteric communication from TMD to NBD regions upon substrate binding. In other words, TMH and NBD motions are not moving altogether as a single block but their motions are coupled, which is assigned to the pivotal role of the TMH-NBD coupling helices as described for type IV ABC transporters ^6,7^. Therefore, PC binding to ^IF^ABCB4 conformation leads to IC opening, which in turn might be associated with the triggering of NBD dimerization.

#### 3.1.2. Impact of the lipid composition onto the overall structural dynamics of ABCB4

Overall structural dynamics in symmetric or asymmetric membranes exhibit similar patterns according to ABC conformational space, even though a larger variability was observed for asymmetric membrane (Figure 1B&C). Similar behaviors were reported for MRP1 MD simulations conducted in different membrane simple models, for which the lipid bilayer was suggested not to significantly affect local minima, but likely kinetics between milestone steps along transport cycle^13^. Taking advantage of our extensive unbiased MD simulations, conformational subspaces were featured in terms of free energy using the InfleCS clustering method^53^ (Figure 2A). It is important to note that in the present study, μs-scale sampling is not sufficient to accurately investigate free energy transitions between conformational states. Likewise, the known TMH kinking upon substrate binding for ABCB4^3^ and ABCB1/P-gp^55^ was not considered here for conformational transition between different states.

**Figure 2.**
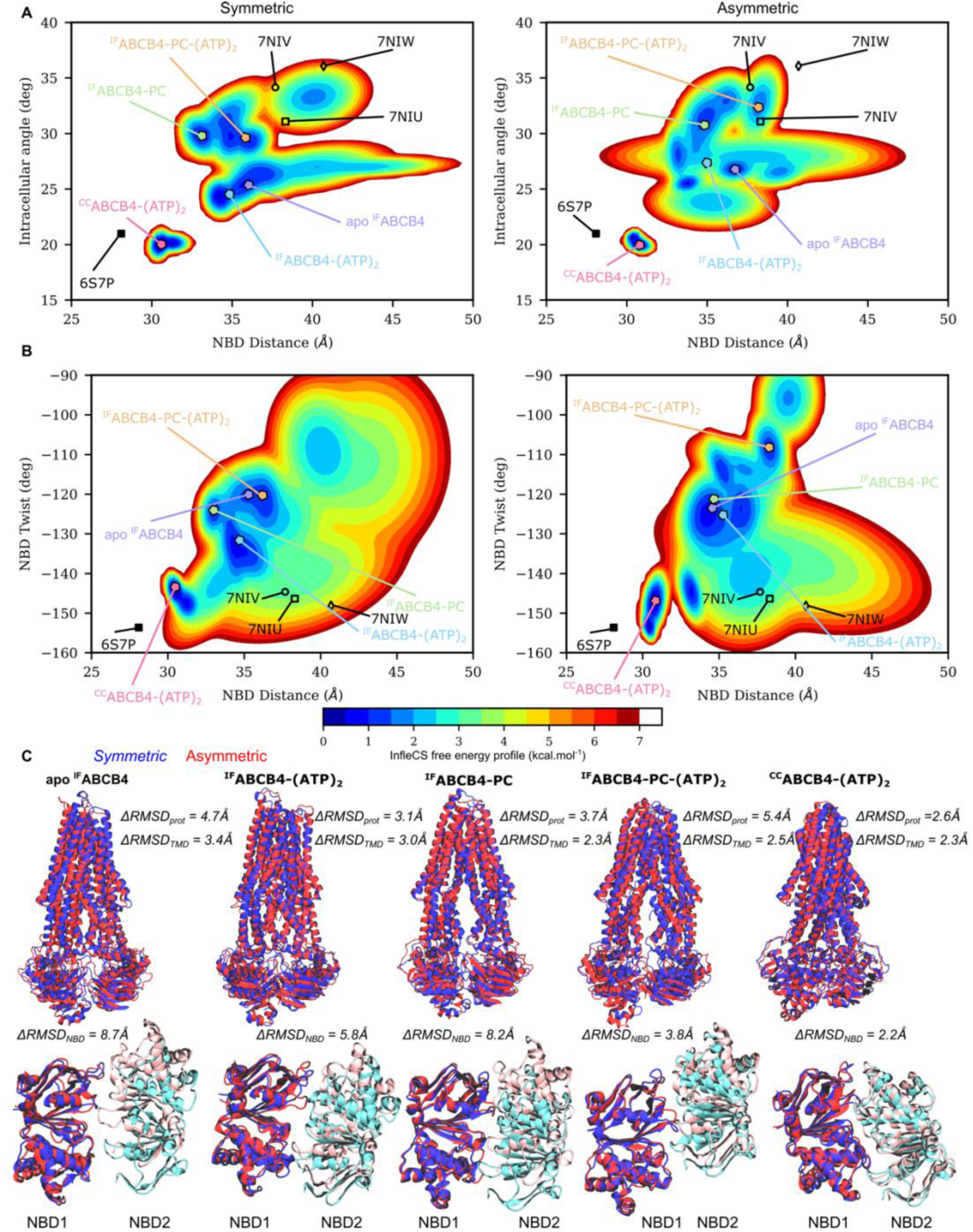
Local conformational sampling of ABCB4 MD simulations. InfleCS free energy-based conformational clustering of ABCB4 under different conformations and bound states in symmetric and asymmetric membrane models according to (A) IC angle versus NBD distance and (B) NBD twist versus NBD distance. (C) Superimposition of representative snapshots for ABCB4 main populations in symmetric (blue) and asymmetric (red) lipid models. NBD1 and NBD2 are respectively depicted in blue and cyan for symmetric lipid bilayer model and red and pink for asymmetric lipid bilayer model. Residues selected for RMSD calculations of all TMH and NBD are reported according to the topology reported in Table S5, the former being made of TMH residues. Protein RMSD calculations were conducted considering the whole protein.

The local free energy surfaces were calculated using the distribution of ABC structural parameters assuming a sufficient sampling of the local conformational space. IC angle, NBD distance and NBD twist were used to monitor free energy surface (Figure 2A&B). We thus identified local minima sampled along MD simulations which may be assigned to metastable states along ABCB4 transport cycle.

We first evaluated different populations per conformation and per bound state (Figures S6-S10) according to the aforementioned ABC structural parameters. MD simulations performed in asymmetric lipid bilayer exhibit a larger structural variability of ABCB4 conformations leading to more conformational sub-states (Table S8 and Figures S6-S10), except for the ^CC^ABCB4- (ATP)_2_ conformation (Figure S10), which is restrained in the same subspace regardless of the membrane composition. Structural overlap between IF states is more favored in an asymmetric membrane than in symmetric lipid bilayer (Figure 1). Therefore, we compiled ABC structural parameters of each conformation and bound state as a single dataset in order to assess potential structural overlaps along alternating access (Figure 2A&B). This is particularly true for ^IF^ABCB4-PC-(ATP)_2_ which strongly overlaps with conformational subspace of ^IF^ABCB4-PC and ^IF^ABCB4-(ATP)_2_ states, even though the main representative snapshot looks to be off. Main representative snapshots obtained from InfleCS clustering in symmetric and asymmetric membranes were superimposed to assess the structural deviations (Figure 2C). While ^IF^ABCB4-PC, ^IF^ABCB4-(ATP)_2_, and ^CC^ABCB4-(ATP)_2_ states exhibit RMSD differences ranging from 2.6 to 3.7 Å, the largest deviations were obtained for apo ^IF^ABCB4 and ^IF^ABCB4-PC-(ATP)_2_. This can be easily explained for the former since apo ^IF^ABCB4 exhibits the largest structural dynamics in both membranes as compared with other states, leading to more clusters. For ^IF^ABCB4-PC-(ATP)_2_ state, we observed that NBDs are not perfectly aligned or anti-symmetrically facing each other (Figure 2C). One NBD appears slightly displaced relative to the other in both membranes. This displacement creates an asymmetry in the NBD arrangement, similar to what has been observed in ^IF^MRP1-LTX- (ATP)_2_.^13^ This shift is more pronounced in asymmetric membrane model, suggesting that conformational transition of the TMD toward OF confirmation not only facilitates PC extrusion but also promotes proper NBD dimerization for ATP hydrolysis.

Since MD simulations demonstrated that similar conformational spaces were sampled regardless of membrane composition (Figures 1 and 2), it can be hypothesized that membrane composition influences conformational transitions rather than significantly altering ABCB4 conformations and structural parameters. Notably, membrane-dependent structural deviations were investigated by focusing on the TMD core and NBD dimer arrangement (Figure 2C). As expected, ^CC^ABCB4-(ATP)_2_ is the only state in which the RMSD values of the TMD core and NBD dimer fall within the same range, reflecting the low structural variability observed during the MD simulations.

#### 3.1.3. Conformation-dependent cholesterol binding spots

We first investigated potential cholesterol binding sites to compare with former investigations conducted on ABCB1/P-gp since it shares 76% identity and 86% similarity with the ABCB4 transporter^14^. Likewise, key cholesterol binding sites were proposed for their cousins MRP1^13^ and ABCB11^15^ suggesting an active role for allosteric communication along the protein. Therefore, we probed membrane- and conformation-dependent cholesterol binding sites of ABCB4 from MD simulations. As observed for P-gp and MRP1, focusing solely on cholesterol recognition consensus motifs (namely CRAC and CARC) is not sufficient to properly describe cholesterol binding. In ABCB4, 4 CRAC and 10 CARC motifs were identified in the transmembrane region, 2 CRAC and 6 CARC motifs for which overlap was observed as for P-gp and MRP1 (Table S9). Cholesterol binding sites were probed by calculating (i) atomistic contacts between cholesterol at the membrane interface along MD simulations and (ii) protein interface for which the likelihood of finding cholesterol molecules is higher than 60% (Figure 4A-4D and Figures S11-S12). Overall, TMH3, TMH4, TMH5, and TMH6 seem to be involved in cholesterol binding spots for both IF and CC conformations. We also observed that a cholesterol hotspot is found between TMH4 and TMH3 for apo ^IF^ABCB4 and ^IF^ABCB4-(ATP)_2_ systems (Figure S11). MD simulations suggested that bound cholesterol may preclude the entry of PC substrate through the TMH4-TMH6 front gate, even though contact analysis showed a smaller contact fraction in the asymmetric membrane (Figure S12). Interestingly, we observed conformation dependence for potential cholesterol binding sites in which two patterns can be identified for apo ^IF^ABCB4/^IF^ABCB4-(ATP)_2_ and ^IF^ABCB4-PC/ ^IF^ABCB4-PC- (ATP)_2_/^CC^ABCB4-(ATP)_2_ (see Figure 4A-D and Figures S11-S12). In PC-bound ^IF^ABCB4 and ^cc^ABCB4-(ATP)_2_ states, additional cholesterol hotspots were observed as shown by numerous atomic contacts between cholesterol and TMH10 and TMH11 (Figure 4A&C and Figure S12). Our results also confirm that CRAC and CARC motifs are not sufficient to probe cholesterol binding for ABC transporters in agreement with observations made for MRP1^13^ and P-gp^14^. Indeed, large scale conformational changes along transport cycle substantially modify residue exposition to lipid bilayer membrane, and therefore protein-lipid interactions. Cholesterol residency time around ABCB4 was calculated highlighting increasing residency time along the transport cycle, i.e., from IF to CC conformation (Figure 4E). This suggests that the surrounding lipids modulate conformational transition of ABCB4.

**Figure 3.**
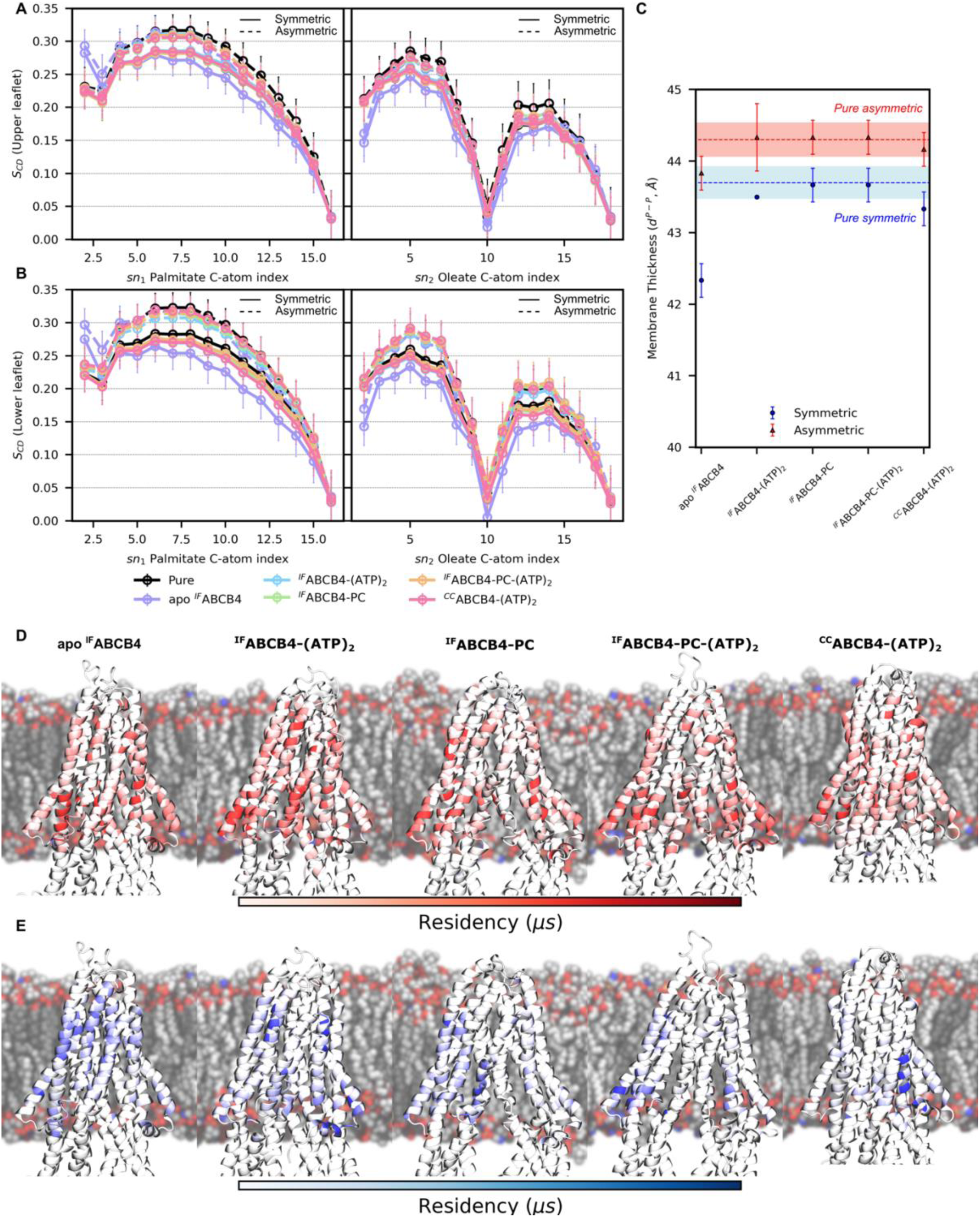
Conformation- and bound-state dependent lipid bilayer properties. Conformation- and bound state-dependent order parameters of palmitate and oleate lipid tails of (A) outer and (B) inner leaflet in symmetric (solid lines) and asymmetric membrane models (dashed line). Pure membrane data were included (black) for sake of comparison. (C) Conformation- and bound state-dependent membrane thicknesses calculated from Phosphate-phostate distances. Pure membrane thicknesses were also shown as dashed lines (shadowed by standard deviations). Calculated residency time for surrounding (D) unsaturated and (E) saturated lipid tails depending on ABCB4 conformational and bound states.

**Figure 4.**
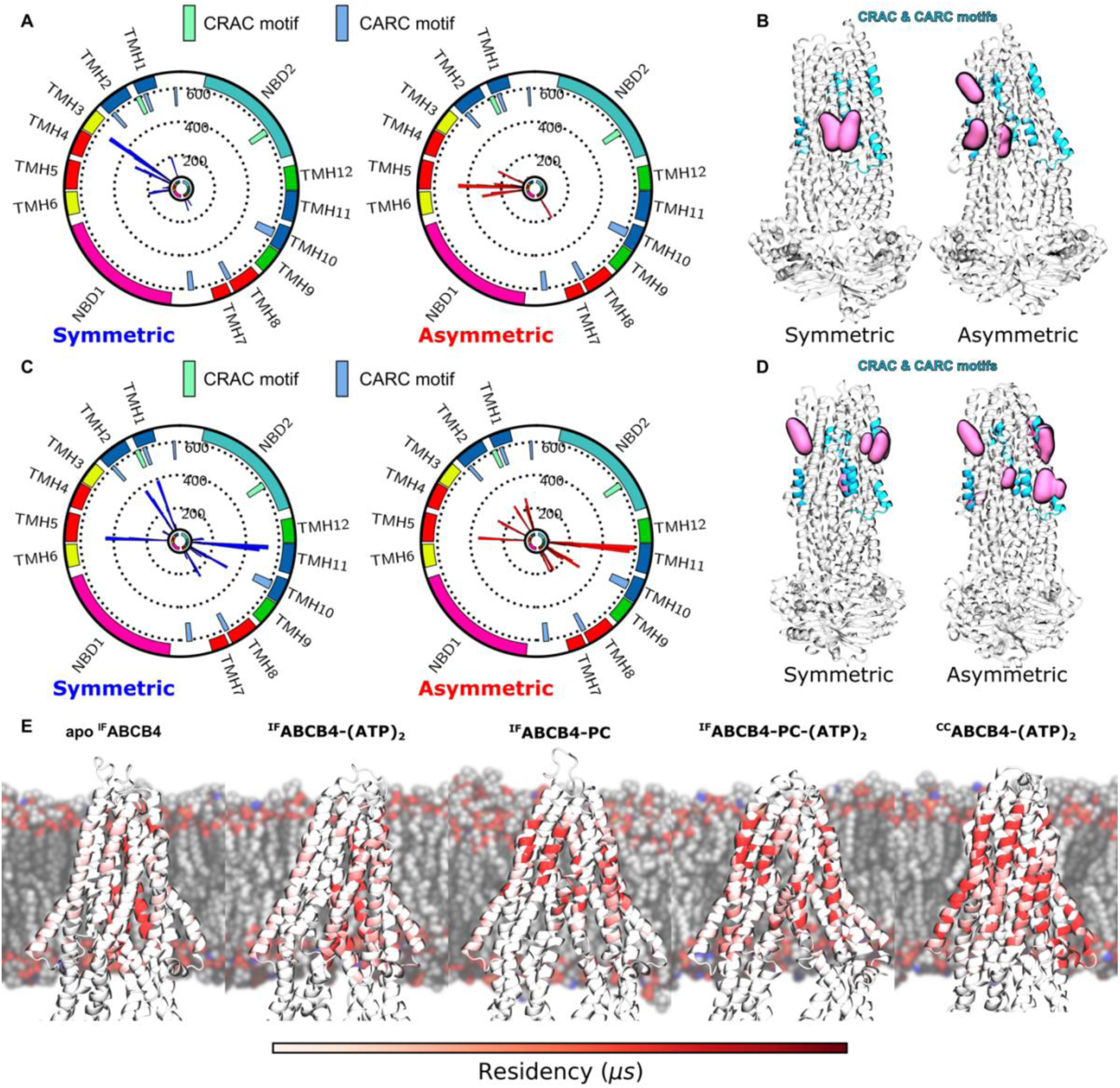
Cholesterol-protein interplay and binding motifs. Conformation-dependent membrane properties Per-residue atomic contacts between cholesterol and (A) apo ^IF^ABCB4 or (C) ^CC^ABCB4- (ATP)_2_. CRAC and CARC motif locations in ABCB4 sequence are respectively depicted in light green and light blue boxes. Calculated cholesterol hotspots (occupancy greater than 60%) for (B) apo ^IF^ABCB4 or (D) ^CC^ABCB4-(ATP)_2_. All CRAC and CARC motif locations are all showed in cyan. (E) Calculated cholesterol residency times depending on ABCB4 conformation and bound-state investigated in this study.

#### 3.1.4. Non-covalent protein-phospholipid interactions

Given the asymmetric repartition of phospholipids, lipid hotspots were mostly observed at the protein-membrane interface in the inner leaflet (Figure 5A). From MD simulations, lipid hotspots for which PC, PE, PS and PA polar heads were present for more than 60% during the simulations were identified and showed in Figure 5A.

**Figure 5.**
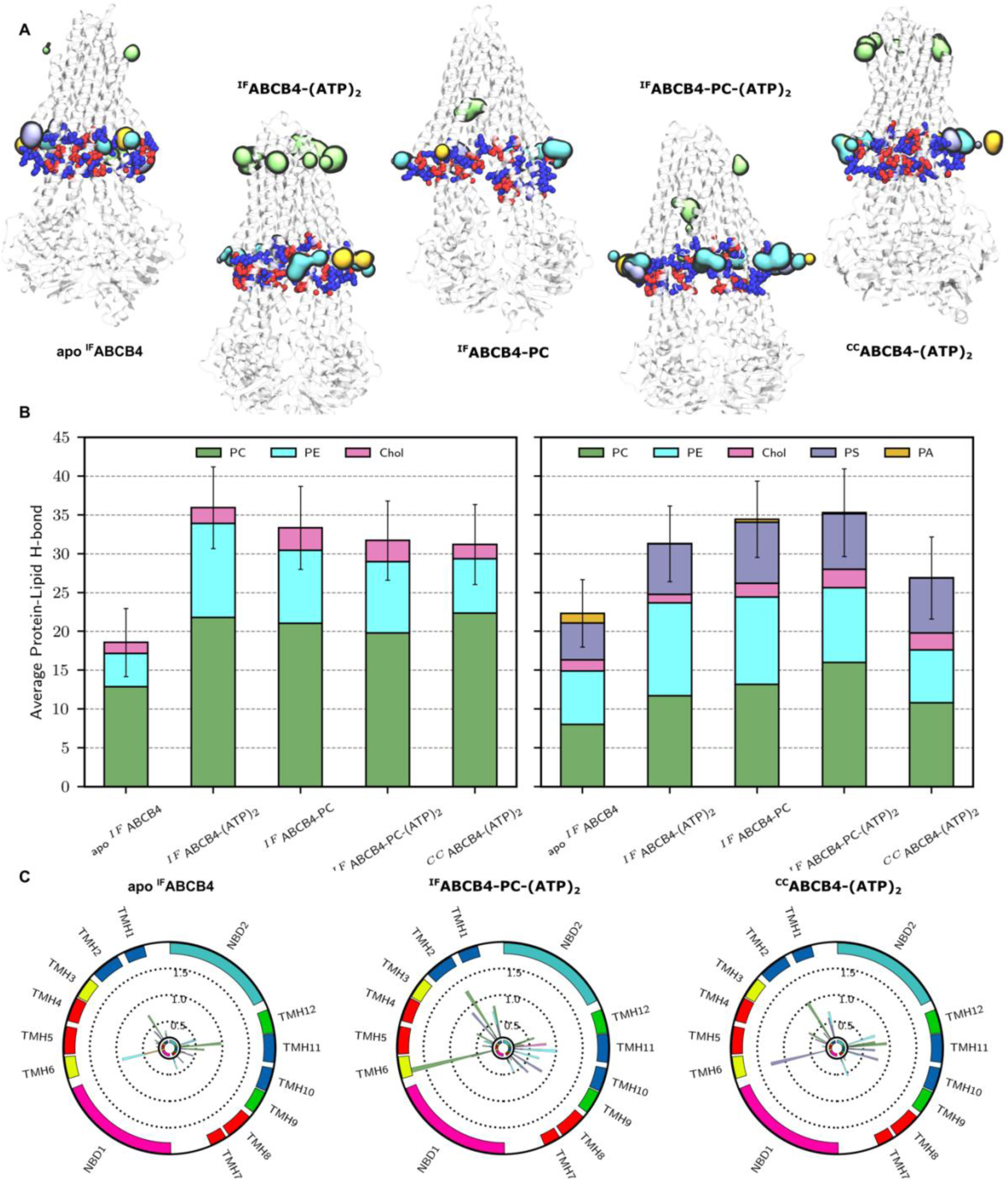
Lipid-protein non-covalent interactions from ABCB4 MD simulations. Conformation and bound state-dependent (A) polar head hotspots and (B) average lipid-protein H-bond counts from MD simulations. (C) Per-domain calculated H-bond fractions over MD simulations for apo ^IF^ABCB4, ^IF^ABCB4-PC-(ATP)_2_ and ^CC^ABCB4-(ATP)_2_.

Interestingly, we observed PS and PA polar heads in close contact with ABCB4 regions rich in cationic residues at the membrane-water interface. Since PS and PA polar heads are negatively charged, this suggests the importance of electrostatic contributions in maintaining polar head lipids in close proximity to ABCB4. Lipid-protein H-bond network was also investigated to rationalize the protein-lipid interplay. Interestingly, H-bond protein-lipid network is globally maintained for both asymmetric and asymmetric membranes. However, apo ^IF^ABCB4 systematically exhibited a weaker H-bond network than all other conformations (Figure 5B). This might be explained by its larger structural flexibility, for which the larger apo ^IF^ABCB4 structural dynamics might lead to more frequent H-bond formation/breaking events. While comparing symmetric and asymmetric membranes, average protein-lipid H-bond counts are similar for all conformations and states but ^CC^ABCB4-(ATP)_2_ for which the average H-bond number is slightly smaller in asymmetric membrane than in symmetric one. In asymmetric membrane, the lower abundance of PC lipids in the inner leaflet is compensated by the presence of anionic PS phospholipids to maintain protein-lipid H-bond network. PA phospholipid poorly participated to H-bond network, suggesting that PA hotspots are mostly driven by electrostatic interactions. However, this should be considered carefully since only ca. 2% of PA lipids are present in asymmetric membrane model. Protein-wise, the overall pattern of protein-lipid H-bond network is slightly affected by membrane composition. While focusing on H-bond only present for more than 35% along MD simulations, 65% of residues are shared between symmetric and asymmetric membranes, with similar scores (Table S10). Overall, the dynamic protein-lipid H-bond network involved mostly intracellular regions of the so-called bundle A, *i.e.*, TMH1, TMH2, TMH11, and TMH12 (see Figure 5C). As expected with the partial atomic charge of polar heads, most of ABCB4 residues involved in protein-lipid H-bond network are polar residues such as glutamate, lysine, and arginine. It is worth mentioning that Glu52 which has been shown to be involved in credit-card swipe mechanism is also strongly involved in protein-lipid H-bond network as its almost neighbor Lys54. Interestingly, these two residues which are respectively anionic and cationic might match with zwitterionic state of PC lipids. MD simulations also revealed the involvement of pre-TMH1 and pre-TMH7 elbow helix residues as key for protein-lipid network (namely, Thr44, Ser696, and to a lesser extend Lys705, see Figure 5C and Table S10). H-bond analyses suggested an almost permanent H-bond with Arg361 located in TMH6, regardless of polar head type. Noteworthy, Table S10 also reported H-bonds breaking from equilibration as well as H-bond that are either formed along MD simulations or strengthen as compared to equilibration stage. Interestingly, pre-TMH1 and pre-TMH7 H-bonds are either formed or strengthened along MD simulations, suggesting a central role of elbow helices in lipid-protein interplay, as observed in MRP1 either from cryo-EM resolution^56^ or MD simulations^13,57^.

### 3.2. PC lipid as substrate of ABCB4

#### 3.2.1. Structural impact of PC substrate canonical binding onto ABCB4 structure

PC binding into the canonical substrate binding pocket led to the opening IC angle associated with the triggering of NBD dimerization when nucleotides are present (Figures 1&2). It is worth mentioning that even if the main population of ^IF^ABCB4-PC-(ATP)_2_ observed from MD simulations exhibited larger NBD distance (See Figure 1), it represents approximately only 43% (populations C1 and C7, see Figure S9).

Other subpopulations (C2 to C6 and C8-C9, see Figure S9) with NBD distances in similar range as ^IF^ABCB4-(ATP)_2_ state correspond to 57% of MD simulations. This might be explained by the cryo-EM structure of PC-bound ABCB4 which is likely more open than *in situ*, and the MD sampling was not sufficient to systematically reach the proper NBD distance. To document on PC substrate entry, we conducted structural analyses of front and back (or rear) entry gates that have been described in the literature for ABCB1 ^55^ and ABCB4^9^. These gates consist of transmembrane regions of TMH3/TMH4/TMH6 or TMH9/TMH10/TMH12 for front and back gates, respectively (Figure 6A). The front gate exhibited a significant structural difference between PC-bound IF conformations and “apo” states (*i.e.*, apo ^IF^ABCB4 and ^IF^ABCB4-(ATP)_2_) as pictured by RMSD (Figure 6B).

**Figure 6.**
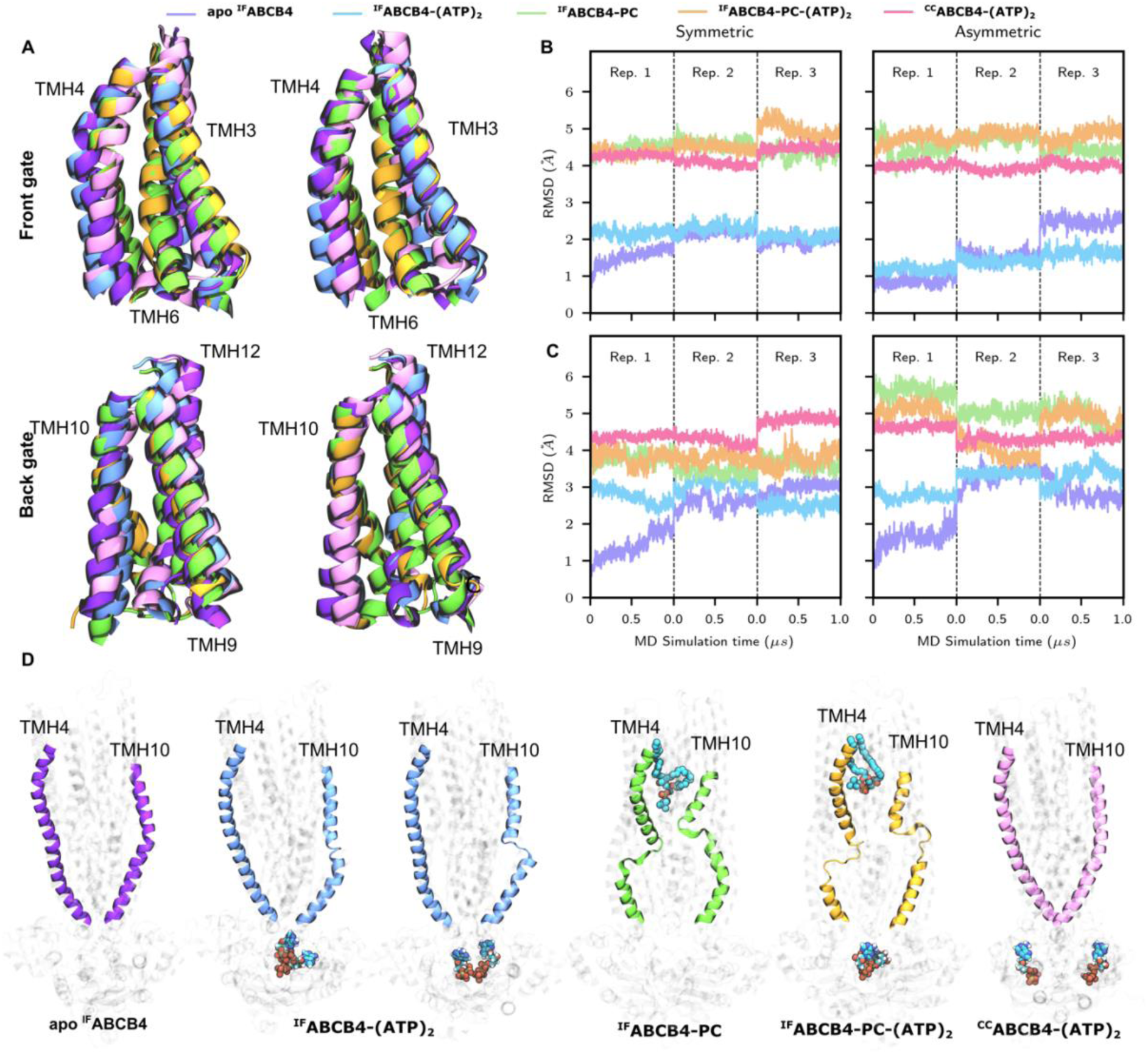
Comparison of TMH arrangement and helicity of ABCB4 conformations and bound states in symmetric and asymmetric membrane. (A) Time-dependent root-mean-squared deviations of ABCB4 front and back gates for all ABCB4 conformations and substrate bound states, using initial frame of apo ^IF^ABCB4 as reference. Superimposition of representative snapshots of (B) front and (C) back gates of ABCB4 conformations and bound states in asymmetric model. RMSD were calculated respectively using Gly189-Leu238 and Ile330-Ala356 as selection for the front gate, and Gln825-Leu878 and Arg969-Tyr997 as selection for the back gate. (D) Conformation- and bound state-dependent TMH4 and TMH10 kinking over ATP and PC binding event along canonical alternating access.

The presence of PC substrate in the canonical binding pocket led to the motion of TMH6 which might be associated with a steric hindrance precluding the entry of other substrates while non PC-bound ^IF^ABCB4 exhibited more open conformation (Figure 6A&B). However, we did not observe any spontaneous PC or other phospholipid entry into the binding pocket. This might be explained by the relatively short sampling of our MD simulations (2μs) while such an event might require longer timescale. To a lesser extent, back/rear gate is more similar between different states but for one replica (Figure 6B&C). MD simulations suggest a TMH10 motion closing the gate entry (Figure 6C&D). ^CC^ABCB4-(ATP)_2_ front and back/rear gates exhibited slightly different conformations as compared to PC-bound states; even though they both remained closed along MD simulations. These observations were made with both symmetric and asymmetric membranes, suggesting that lipid composition does not have a relevant impact.

We performed PCA analysis considering each conformation and each state to better decipher the impact of PC substrate binding onto the TMH arrangement of ABCB4 (See Figure S13). Both TMH4 and TMH10 were identified as main source of structural variability along the alternating access. In presence of PC substrate, TMH4 and TMH10 exhibited kinking event leading to an important discontinuity in TMH helicities (Figure 6D and Table S11). In other words, the presence of PC substrate was associated with TMH4 and TMH10 kinking and adopting a Y-shaped cavity^3^ likely to restrict substrate in the binding pocket, precluding its exit. Our findings are in agreement with observations made with ABCB1/P-gp^16^. This was also confirmed by measuring TMH helicity along MD simulations (Table S11), in which *e.g.*, TMH4 and TMH10 exhibit helicities of 0.94 ± 0.21 and 0.85 ± 0.25 in apo ^IF^ABCB4 and 0.78 ± 0.36 and 0.69 ± 0.35 in ^IF^ABCB4-PC-(ATP)_2_ in asymmetric membrane.

Interestingly, kinking of TMH10 was also observed in at least one replica of ^IF^ABCB4-(ATP)_2_. Noteworthy, both TMH4 and TMH10 are directly involved in the interactions between NBDs and TMDs through the “ball-and-socket” arrangement between NBDs and TMH through the intracellular coupling helices including those connecting TMH4-TMH5 and TMH10-TMH11. Therefore, proper ATP binding might propagate to TMH domains as proposed for other ABC transporters^10,58^. One can thus assume a synergistic effect of ATP and PC substrate binding to favor such an event, even though the latter might be sufficient.

#### 3.2.2. Structural impact of PC substrate canonical binding onto ABCB4 structure

Taking advantage of the cryo-EM structure of ^IF^ABCB4-PC state, we investigated key amino acid involved in PC binding (Figure 7A). In line with cryo-EM structure, the central role of Trp234, Phe345, His989^3^ (located in TMH4, TMH6, and TMH12, respectively) were also observed in our MD simulations. For instance, calculated atomic contact fractions in asymmetric model are respectively 9.23, 30.64 and 25.63 for ^IF^ABCB4-PC and 29.42, 26.52 and 19.63 for ^IF^ABCB4-PC-(ATP)_2_ (see Table S12). MD simulations also suggested Phe305 (TMH5) and Gln725 (TMH7) as involved in PC substrate binding (Figure 7A), contact fractions being respectively 17.14 and 22.38 for ^IF^ABCB4-PC and 4.61 and 22.15 for ^IF^ABCB4-PC- (ATP)_2_. The presence of three aromatic residues suggests a network of cation-π dispersive interactions maintaining PC substrate into a so-called cation-π cage (Figure 7A).

**Figure 7.**
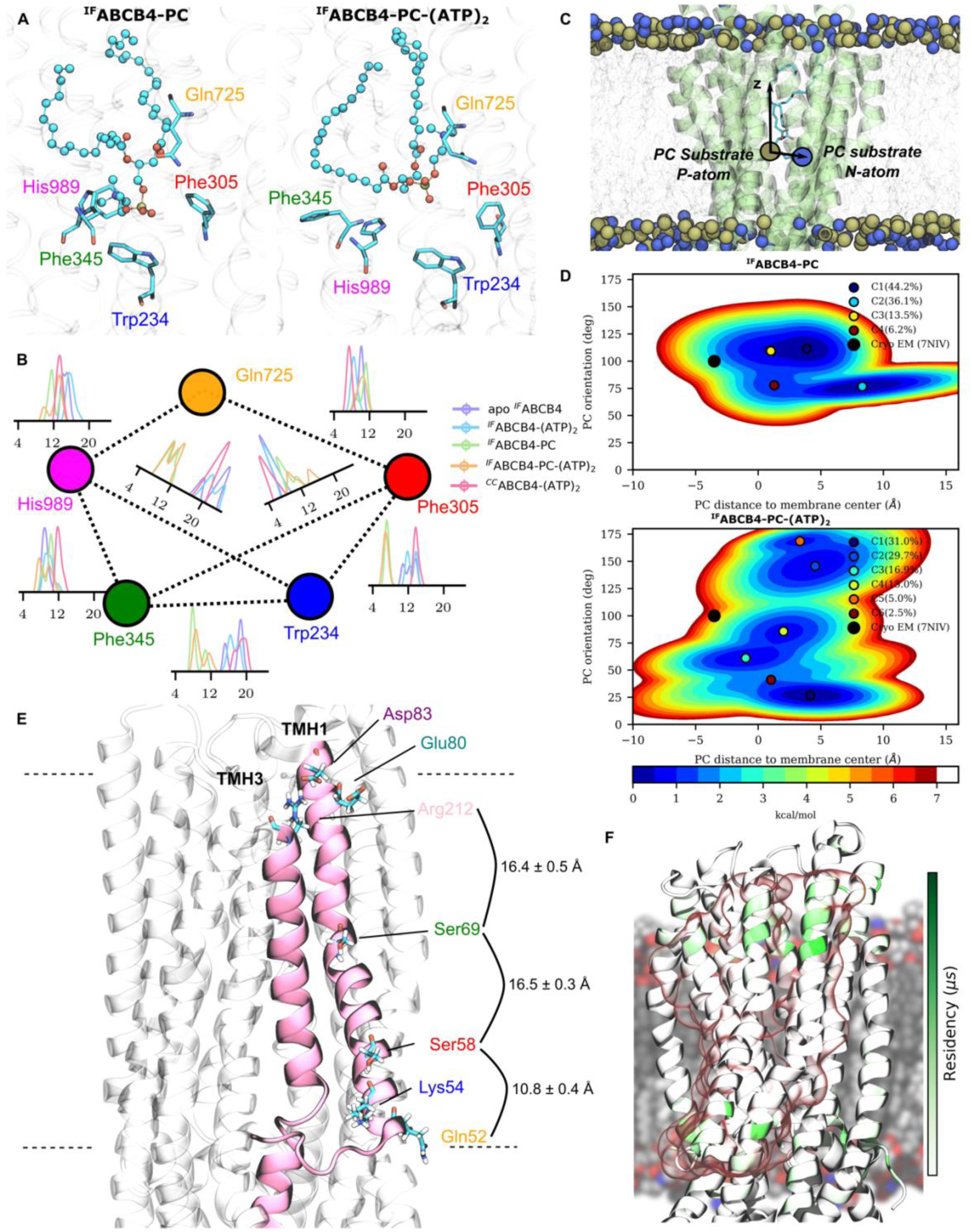
PC substrate binding in canonical and credit-card swipe alternating access. (A) Representative snapshots of ^IF^ABCB4-PC and ^IF^ABCB4-PC-(ATP)_2_ in asymmetric model highlighting identified residues involved in cation-π cage and (B) conformation and bound-state dependent inter residue distances of the binding pocket network. (C) Definition of geometrical parameters, namely PC polar head orientation and distance to membrane center used to assess (D) the local free energy surface of substrate binding event of canonical alternating access. (E) Mapping of potential polar amino acid involved in credit-card swipe mechanism including key average distances along ^CC^ABCB4-(ATP)_2_ TMH1 and TMH3 and (F) calculated residency time of PC polar head for TMH1, TMH2 and TMH3 of ^CC^ABCB4-(ATP)_2_ from MD simulations.

Noteworthy, Trp234 was also proposed to play a central role in (i) substrate entry and then (ii) maintaining PC substrate in the canonical hydrophobic binding pocket by means of cation-π interactions^3^. His989 and Gln725 play also an important role through H-bond network between phosphate moiety with residue side chains (Table S13). For instance, our MD simulations showed that PC substrate and His989 side chain were involved in H-bond for 35% (26%) and 32% (6%) respectively for ^IF^ABCB4-PC-(ATP)_2_ and ^IF^ABCB4-PC in asymmetric (symmetric membrane). Likewise, Gln725 and PC substrate interact by H-bond for 15% (27%) and 7% (33%) for ^IF^ABCB4-PC-(ATP)_2_ and ^IF^ABCB4-PC in asymmetric (symmetric membrane), respectively. We can observe that H-bond exchange might occur between His989 to Gln725 (Figure 7A), the former being deeper in membrane than the latter. Assuming that His989, Gln725, Phe305, Phe345 and Trp234 form a binding network, we calculated conformation-dependent inter-residues distances (Figure 7B). PC binding is systematically associated with shorter Trp234-Phe305, Trp234-Phe345 and Trp234-His989 distances and slightly larger for Phe305-Phe345. Distances between these residues and PC polar head were also monitored (Figure S14) confirming the non-covalent network driven by H-bond and cation-π interactions. Interestingly, the presence of ATP in NBSs is associated with slightly larger distances between PC polar head and aromatic residues. Taking advantage of our 1 µs MD simulations, we also estimated the free energy surface of PC substrate according to the distance of PC polar head with membrane center and the orientation of PC polar head as compared to membrane normal axis (Figure 7C&D). We considered ^IF^ABCB4-PC-(ATP)_2_ and ^IF^ABCB4-PC to assess the coupling between nucleotide binding to NBS with PC substrate (Figure 7A). Even though the sampling of our MD simulations was not sufficient to observe expected PC flip-flop in the binding pocket, the presence of ATP bound to both NBS led to a local free energy minimum in the conformational space of PC polar head crossing the membrane center and properly oriented towards outer leaflet (Figure 7D).

Noteworthy, in absence of ATP, PC polar head never reoriented itself toward outer leaflet, even if most of local minima have crossed membrane center. One can thus assume that proper PC binding is sufficient to help PC polar head to cross membrane center, but ATP binding and subsequent TMH arrangement are required to favor PC polar head flip-flop.

#### 3.2.3. Structural insights into the potential credit-card swipe mechanism from μs-scale MD simulations

Credit-card swipe PC translocation was described from non-conserved polar amino acids (Gln52, Ser58 and Ser69) between ABCB1 and ABCB4, the former being a “drug transporter” for which PC translocation has not been formerly confirmed yet ^16,59,60^. These amino acid side chains are oriented toward the membrane in ^CC^ABCB4-(ATP)_2_ conformation. Our μs-scaled MD simulations are in agreement with previous observations made from either shorter unbiased simulations (200 ns, 5 replicas) or potential of mean force calculations^9^. Side chain orientations toward lipid bilayer membrane were maintained, even for the deep Ser69 which is not located at the interface between high-density polar head and high-density lipid tail regions (Figure 7E). In line with predicted PC polar head hotspots, MD simulations also suggested longer PC residency times TMH1-3 residues at the water-membrane interface, suggesting preferential non-covalent interactions along the credit-card swipe pathways (Figure 7F). Unfortunately, we were not able to observe PC flip-flop events along our MD simulations nor specific binding to those amino acids, which is in contrast to the previous work of Prescher et *al.*^9^. This could be explained by the (i) insufficient time scale from our μs-MD simulations and (ii) the estimated free energy barrier obtained by umbrella sampling simulations ranging from 10 and 20 kcal.mol^-1^.

## 4. Discussion

In the present study, all-atom MD simulations were carried out to investigate the overall conformational space and the lipid-protein interplay of ABCB4. Several conformations and bound states were considered taking advantage of cryo-EM structures resolved in nanodiscs mimicking a native environment of the canalicular membrane. By conducting μs-long unbiased MD simulations, ABC structural patterns of ABCB4 conformations and bound states were investigated considering two lipid bilayer membrane models, namely symmetric (made of POPC:POPE:Chol (2:1:1)) and asymmetric. Protein-lipid interplay was investigated by assessing (i) the impact of membrane composition on the structural dynamics of ABCB4 and (ii) conformation-dependent protein-lipid contact patterns driven by non-covalent interactions. Special attention was also made on substrate-ABCB4 interactions considering either a canonical alternating access^3^ as well as a credit-card swipe mechanism^9^. In line with recent observations made for ABCB4 cousin, namely P-gp/ABCB1^16^, TMH4 and TMH10 kinking was also investigated given the different TMH conformations observed from cryo-EM resolved structures. Large-scale conformational transitions were not investigated in the present study given the challenge to define the proper driving force responsible for either IF-to-OF or OF-to-UR conformational transitions.

### 4.1. Membrane-dependent structural dynamics of ABCB4

While ABCB1/P-gp has been extensively studied over the past decades, recent studies highlighted that knowledge of ABCB family still requires further investigations. This is particularly true for ABCB4 which is not a drug transporter even though studies showed that few xenobiotics may be translocated by ABCB4^9,61,62^. It is important to note that the present study is limited by forcefield inaccuracies as well as limited conformational sampling. However, our set of MD simulations appear relevant to decipher local conformational space of each conformer and bound state including potential overlaps.

In contrast to what has been observed from MD simulations of ABCC1/MRP1^13^, ABC structural parameters (*i.e.*, IC angle, NBD distance, and NBD twist) measured along the ABCB4 simulations remain close to cryo-EM experimental structures. This can be easily explained by MRP1 cryo-EM resolution in detergent leading to wide open IF structure. One can also hypothesize that the degenerated NBS in C-family ABC transporter can be associated with permanent single ATP-bound state requiring the closing of IC gate and NBD dimerization^13^. Notably, ATP hydrolysis for non-degenerated ABC transporter as ABCB4 was however proposed to alternate single ATP hydrolysis from NBS to another along several transport cycles ^63^.

Our MD simulations are in line with previous experimental and theoretical investigations for which ABC IC angle and NBD distance motions are not necessarily coupled. This is particularly true for ABCB4 while considering the canonical alternating access since PC substrate binding is associated with increase of IC angle owing to steric hindrance in the binding pocket. On the other hand, NBD distance shortening requires nucleotide binding since ATP is known to be sandwiched by both NBDs^6^. However, none of our MD simulations conducted with IF ATP-bound state (*i.e.*, ^IF^ABCB4-(ATP)_2_ and ^IF^ABCB4-PC-(ATP)_2_) reached proper NBD dimerization as pictured by NBD twist distribution. This is due to the insufficient sampling at the μs-scale. One can infer that proper ATP hydrolysis competent NBD dimerization, as observed in ^CC^ABCB4-(ATP)_2_, requires TMD large scale conformational transition which then propagate to NBD conformation. We observed translation of NBDs along the membrane when bound to both PC substrate and ATP (*i.e.*, ^IF^ABCB4-PC-(ATP)_2_). Interestingly, similar shifted NBD arrangement was also observed for MRP1 bound to nucleotides and substrate ^13^. One can hypothesize from these MD simulations conducted with two mammalian ABC transporters that such NBD conformation is a transient state prior to TMD large scale conformational transition. Unfortunately, there is not a single resolved cryo-EM structure of mammalian fold IV ABC transporter to validate such hypothesis.

The MD simulations did not reveal substantial impact of membrane composition onto conformational local minima. Overall, the same conformational space was sampled along MD simulations regardless of membrane composition. Interestingly, asymmetric membrane simulations showed more structural variability and flexibility as compared to simulations conducted in symmetric membrane. The larger structural variability in asymmetric membrane and proposed lower energy barriers between different ABC conformations suggest that membrane composition might mostly affect kinetics of ABCB4 transport cycle. However, free energy surfaces proposed in the present manuscript must be deemed very carefully in the framework of alternating access. ABC structural parameters can only be considered to monitor large-scale conformational transitions for type IV ABC transporters. However, defining driving forces responsible for conformational changes remains an important challenge in ABC transporter’s alternating access. Recent experimental investigations conducted on MRP4 have proposed that lipid environment might modulate, at least ATPase activity^17^, for which native environment was associated with higher activity than in POPC-reconstituted nanodiscs. In other words, lipid composition is likely important for ABC transporter kinetics.

### 4.2. Protein-lipid interplay and annular lipids of ABCB4

Present MD simulations were also used to investigate protein-lipid interactions and individuate potential conformation-dependent lipid fingerprints. First, by monitoring membrane properties, we observed that sn_1_-palmitate and sn_2_-oleate lipid order parameters were systematically more ordered in asymmetric membrane than in symmetric model. Such a finding is counter-intuitive with the larger flexibility observed for asymmetric conformational sampling in asymmetric membrane. Noteworthy, lipid order parameters were calculated only for sn_1_-palmitate and sn_2_-oleate lipids for sake of comparison with symmetric membrane. Therefore, ∼20% of lipid tails were not included to properly assess membrane fluidity. Besides, membrane thicknesses were calculated for which asymmetric membrane is ∼2 Å thicker than symmetric one, leading to larger protein-lipid interface. Altogether, these findings including those regarding ABCB4 conformational flexibility raise the question of the iterative interplay between protein and membrane lipid bilayer, *i.e.*, how do lipids modulate protein flexibility as well as in turn how protein affect membrane properties. Interestingly, we observed that unsaturated lipids exhibit a longer residency time around ABCB4 than saturated lipid tails. Even though these results should be considered carefully given the insufficient timescale to capture lateral diffusion, they suggest that unsaturated lipid tails (*e.g.*, oleyl and arachidonyl) may create a disordered pocket around the protein, which might in turn increase local conformational flexibility and thus the increased protein dynamics in asymmetric membrane. However, this effect might be counterbalanced along the transport cycle. Indeed, the ^CC^ABCB4 pocket is likely to be stiffened owing to its enriched environment in cholesterol as compared with *e.g.*, apo ^IF^ABCB4 and ^IF^ABCB4-(ATP)_2_. One can assume that the more disordered pocket around IF conformation might favor slight structural flexibility required for PC substrate entry.

Our results highlighted conformational-dependent pattern of lipid protein contact and non-covalent interactions. While it is known that the use of CRAC and CARC motifs, at least in ABC transporters ^13,14^, is not sufficient to predict and investigate cholesterol binding, MD simulations clearly suggest that cholesterol binding as allosteric partners differs from one conformation to another (*e.g.*, ^IF^ABCB4 and ^CC^ABCB4). Interestingly, as observed in MRP1^13,57^, we observed cholesterol hotspot in the pre-TMH elbow helix suggesting a specific role of cholesterol in this region. Given that annular lipids, including cholesterol, were shown to actively participate to distant communication between *e.g.*, NBS and substrate binding site, one can hypothesize that cholesterol binding might play different roles in IF-to-OF transition and OF-to-UR-to-IF transition. As it has been very recently suggested for P-gp^16^, protein-lipid interplay in the inner leaflet remains more important than the outer leaflet. It is important to note that this has been suggested only for ^IF^P-gp. Therefore, one cannot conclude regarding CC conformations. However, in the present asymmetric model, anionic lipids are exclusively located in the inner leaflet. Strong electrostatic interactions between anionic polar head and cationic ABCB4 residues are suspected to maintain strong non-covalent network which also differs from one conformation to another. This may be explained by different patterns in terms of residue exposure to high density polar head region according to TMH arrangement. Even though the allosteric role of annular lipid was not within the scope of the present study, one can infer from other studies^13–18^ that membrane itself is not only a physical container for ABCB4 but annular lipids are active player for its function. We can thus expect that lipid composition plays a role in modulating ABCB4 function given that cell membrane structure is highly dynamic and for which, *e.g.*, nanodomains and/or lipid rafts might also affect protein structure and dynamics.

However, our present results regarding the protein-lipid contact should be considered very carefully since only all-atom MD simulations were conducted. In such an approach, the sampling and force field inaccuracy leads to a poor description of lipid lateral diffusion. On the contrary, in the recent study regarding P-gp^16^, coarse-grained model was used with Martini 3 force field which was shown to provide robust insights regarding surrounding lipid fingerprint. Besides, it is worth mentioning that all atom MD simulations using asymmetric membranes remain challenging and should be considered carefully. Indeed, asymmetric membrane is associated with differential lateral compressibility between inner and outer leaflets. Nevertheless, the present study only paves the way for further investigations.

### 4.3. TMD-NBD coupling improve canonical alternating access substrate translocation

Particular attention was paid to substrate entry and interactions in the canonical binding pocket. In contrast to MRP1, P-gp and ABCB4 were shown to bind substrate directly from lipid bilayer, through either the front or rear gate made of TMH3/TMH4/TMH6 or TMH9/TMH10/TMH12, respectively ^55,56,64^. No substrate entry was observed in our simulations with either apo ^IF^ABCB4 nor ^IF^ABCB4-(ATP)_2_. This might be either due to insufficient sampling or the so-called cholesterol gating as proposed for ABCG2. Indeed, our MD simulations showed a preferential cholesterol binding site in this region^65^. However, our simulations clearly demonstrate two distinct supramolecular arrangements of the front gate. On one hand, apo ^IF^ABCB4 and ^IF^ABCB4-(ATP)_2_ exhibit an open front gate with space between TMH3 and TMH4. One the other hand, ^CC^ABCB4-(ATP)_2_ conformation and PC-bound ^IF^ABCB4 states showed TMH6 shift between TMH3 and TMH4. We can here hypothesize that PC substrate binding leads to TMH6 motions closing the front gate in order to preclude (i) substrate exit back to the inner leaflet and (ii) other PC lipid entry. Cryo-EM structures of ^IF^ABCB4 highlighted the central role of TMH4 Trp234 residue as PC substrate carrier through cation-π interactions between cationic choline moiety and aromatic tryptophane benzimidazole group. It is important to note that current force fields tend to underestimate such dispersive interactions^66^, which can explain why we were not able to observe such events. Despite, MD simulations strengthen the proposed central role of cation-π interactions in the substrate binding pocket. Our results confirmed the central role of Trp234 (TMH4), Phe345 (TMH6) and His989 (TMH12) in maintaining PC substrate as shown by mutagenesis experiments. Furthermore, structural dynamics of ABCB4 also identified two other residues, namely Phe305 (TMH5) and Gln725 (TMH7) which might be involved in PC substrate binding and dynamics. The former is involved in maintaining the cation-π cage between aromatic ring and choline cationic moiety while the latter is proposed to favor PC polar head translocation across membrane center given its position along membrane normal. While comparing substrate position with MD simulations, PC polar head spontaneously crossed the membrane center. MD simulations highlighted the already known distant coupling between nucleotide binding and substrate binding pocket since PC polar head tends to flop only in presence of ATP bound to NBS. This might suggest that PC substrate may reorient itself along transport cycle. However, the full flopping event was not observed owing to the present insufficient sampling, while being within the current standard for unbiased all-atom MD simulations. Interestingly, PC flopping was not initiated in symmetric membrane. It could be explained by the insufficient sampling especially for such a POPC:POPE:Chol (2:1:1) symmetric model. Indeed, this is in line with the aforementioned lower ATPase activity of MRP4 observed in POPC as compared to native nanodiscs^17^. Therefore, this suggests that asymmetry might also be an important driving force for substrate translocation by ABC transporters.

In contrast to C-family ABC transporters, ABCB1 and ABCB4 were resolved with kinked TMH4 and TMH10 helices in presence of substrate. TMH kinking is expected to at least maintain ABCB4 substrate in substrate binding pocket. The role of membrane and annular lipids in modulating the conformational space and flexibility of P-gp TMHs was thoroughly investigated. Our MD simulations also suggest that TMH4 and TMH10 flexibility may participate in substrate translocation by carrying PC lipid to the outer lipid-water interface. Our MD simulations carried out with ABCB4 are in agreement with P-gp MD simulations^16^, exhibiting TMH4 and TMH10 kinking events in presence of substrate. Noteworthy, this can be associated with high identity (similarity) scores between ABCB4 and P-gp TMH4 and TMH10, being 85% (93%) and 76% (93%), respectively. However, we did not observe spontaneous kinking events in apo ^IF^ABCB4 systems in spite of longer MD simulations. Spontaneous kinking of TMH10 in absence of PC substrate was only observed when ATP is bound to NBS highlighting the coupling existing between NBDs and TMDs. This can be easily explained by the “ball-and-socket” arrangement between coupling helices and NBD^6^. This is particularly true since TMH4 and TMH10 are directly connected to two of four coupling helices. We can thus assume that proper ATP binding may favor TMH4 and TMH10 kinking event. It seems however that TMH4 and TMH10 kinking remains a specific event for at least few members of B-family ABC transporters. This can also be correlated to the existence of two canonical ATP-hydrolysis competent NBS as compared to *e.g.*, C-family ABC transporters for which no TMH kinking event was reported so far, even in presence of substrate and in native environment. Furthermore, ^CC^ABCB4-(ATP)_2_ exhibit straight TMHs clearly suggesting that substrate translocation might be associated to spontaneous unkinking event which does not require ATP hydrolysis.

### 4.4. Credit-card swipe mechanism as complementary to potentiate ABCB4 PC floppase activity

Another translocation mechanism has also been proposed regarding PC flopping mediated by ABCB4, supported by joint experimental and computational investigations ^9^. The so-called “credit-card swipe mechanism” relies on TMH-mediated translocation events in which a PC lipid located in the inner leaflet PC is expected to bind to polar residues exposed at the protein-membrane interface, namely Gln52, Ser58, and Ser69^9^. Importantly, biased MD simulations were carried out strengthening this hypothesis supported by site-directed mutagenesis experiments demonstrating that mutations of these residues are associated with lower ATPase activity^9^. We monitored amino acids that might favor proper flip-flop of PC substrate (*i.e.*, lipid tail reorientation, see Figure 7E&F). We identified polar residues in the outer leaflet exhibiting side chain orientation toward lipid tails. They may participate in shielding zwitterionic PC polar head through electrostatic and H-bond interactions, namely Arg212 (TMH3), Glu80 (TMH1), and Asp83 (TMH1). Average distance between Arg212 and Ser69 were measured exhibiting similar values as between Ser58 and Ser69 (ca. 16Å).

Likewise, calculated densities (Figure 6E) exhibited Arg212-Ser69 distance ranges from 14.4 and 18.2 Å in asymmetric membrane (Figure 6D). Noteworthy, PC polar head cross sectional diameter was estimated at ca 10 Å^67^. This suggests that while bound to Ser69, PC is within the range of short-distance electrostatic interactions, and translocation may happen from Ser69 to Arg212. In other words, even though we could not confirm PC translocation from all atom MD simulations owing to too short time scale, our results do not discard former investigations suggesting that such a mechanism remains a potential event with ABCB4. Indeed, present MD simulations clearly showed strong H-bond interactions involving Gln52 and Lys54 with surrounding lipids, including PC lipids in spite of their lower abundance in the inner leaflet. Interestingly, by monitoring the different polar residues which may be involved in carrying PC lipid translocation from inner and outer leaflet, we identified a network of polar residues at the outer lipid-water interface, namely Arg212, Glu80 and Asp83. These residues tend to form a dynamic salt-bridge network in the high density polar head region of membrane outer leaflet. This was also confirmed by calculated PC polar head hotspots as well as longer residency time for PC lipids at the water-membrane interfaces of TMH1, TMH2 and TMH3. Our results still require to be confirmed by further investigations, *e.g.*, by (i) considering coarse grained MD simulations to reach sufficient timescale of observing TMH1-assisted PC translocation or (ii) site-directed mutagenesis experiments by mutating Arg212, Asp83, and Glu80. Furthermore, previous work reported that TMH-assisted PC translocation is a very slow event owing to its high activation barrier ranging from 10 to 20 kcal.mol^-1^ ^9^. Therefore, we can hypothesize that credit-card swipe mechanism may occur only if TMH-assisted PC translocation is associated with the CC-to-UR large-scale conformational changes.

Overall, assuming both the canonical alternating access and credit-card swipe mechanisms, we can hypothesize that ABCB4 might be more efficient to favor PC translocation from the inner leaflet to the outer leaflet. Indeed, by mutating key PC residues of canonical alternating access, PC translocation was shown to be less efficient but still possible (from ∼ 20 to 50% of wild-type ABCB4 activity when mutating Trp234, Phe345, and His989). This may suggest that the credit-card swipe mechanism can be complementary translocation event for ABCB4. Assuming TMH1 and TMH3 are involved in TMH-assisted PC translocation event while ABCB4 adopt the CC conformation, one can consider that at least two PC flop events can occur per ABCB4 transport cycle, during either IF-to-OF or CC-to-IF transitions, as pictured in Figure 8. Obviously, such a hypothesis requires further experimental evidence to be carried out.

**Figure 8.**
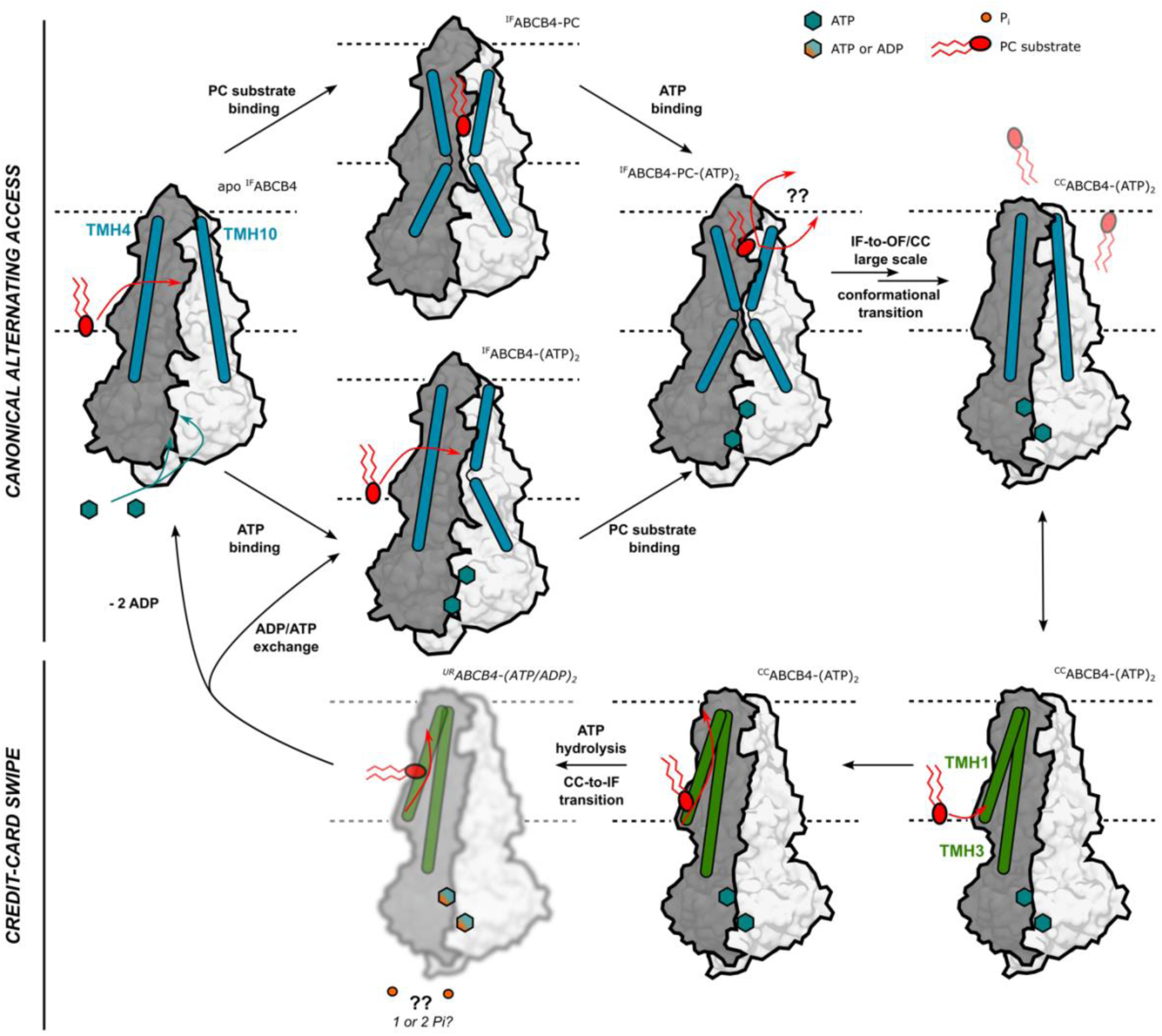
Proposed overall dual mechanism for PC translocation. Canonical alternating access is shown, highlighting TMH kink triggered by either PC substrate and ATP binding. Cryo-EM resolution of ^cc^ABCB4-(ATP)_2_ as well as present MD simulations suggest that TMH helicity is restored during the IF- to-OF/CC conformational transition. Potential credit-card swipe mechanism is also shown in which PC lipid may enrich outer leaflet through a TMH-assisted translocation event. It remains unknown if TMH-assisted translocation event requires CC-to-IF conformational transition. Since ABCB4 has not been resolved under turnover conditions, the ^UR^ABCB4-(ATP/ADP) conformation is blurred.

## 5. Conclusion

Altogether, the present investigation relying on μs-scaled MD simulations provides robust hints regarding the ABCB4 function by conciliating both canonical alternating access and credit-card mechanisms as crucial for ABCB4-mediated PC transport. ABCB4 is not referred to as drug membrane transporter but as an essential physiological transporter to maintain bile composition and homeostasis. Therefore, the selectivity and specificity of ABCB4 dysfunction cannot be overcome by increasing functions of other transporters as shown for membrane drug transporters (*e.g.*, MRP2/MRP4 pair). This is even more important for ABCB4 which acts only in the liver canalicular membrane while drug transporter compensation can exist remotely according to the remote sensing signaling theory. We also strengthen the importance of surrounding lipids and membrane for modulating structural features that have been proposed for ABCB4 cousin, namely ABCB1, such as TMH4 and TMH10 kinking. These results require further investigations to be also considered for other mammalians B-family ABC transporters, in contrast to C-family ABC transporters for which the transport cycle might differ.

## Supporting information

Electronic Supplemental Information

## 6. Data Availability

Initial and final structures for all simulations, MD input examples, ATP force field parameters used, MD trajectories, analysis scripts are available upon reasonable request.

## 7. Acknowledgments

The authors gratefully acknowledge support from Xavier Montagutelli (University de Limoges, France) for computational and technical support, from the regional supercomputers CALI (“CAlcul en Limousin”) and “Baba Yaga”, as well as from the “Jean-Zay” national^56^ supercomputer from IDRIS HPC resources under the allocations 2020-A0080711487, 2021-A0100711487 and 2022-A0120711487 made by GENCI. We also thank Prof. Lutz Schmitt for fruitful scientific discussion and suggestions about the present manuscript.

## 8. Author Contributions

V. Crespi: Conceptualization, Investigation, Methodology, Data curation, Formal analysis, Writing – Original draft, Writing – review & editing

A. Tóth: Methodology, Formal analysis, Writing – review & editing

A. Janaszkiewicz: Methodology, Formal analysis, Writing – review & editing

T. Falguières: Investigation, Supervision, Project administration, Writing - review and editing

F. Di Meo: Conceptualization, Investigation, Methodology, Data curation, Formal analysis, Funding acquisition, Project administration, Supervision, Writing – original draft, Writing - review and editing

## 10. Declaration of generative AI and AI-assisted technologies in the writing process

During the preparation of this work, the authors used ChatGPT 3.5 in order to check grammar and spelling as well as to improve readability of the manuscript. After using this tool, the authors reviewed and edited the content as needed. The authors take full responsibility for the content of the published article.

