## Supplementary material for "Membrane-Dependent Dynamics and Dual Translocation Mechanisms of ABCB4: Insights from Molecular Dynamics Simulations": Electronic Supplemental Information

### List of Supporting Tables

### Supporting Tables

**Supporting Table S1. Lipid composition (number of lipids) for symmetric membrane models.**

| Lipids | Full name (Lipid21 residue topology) | Leaflet | apo $^{IF}$ABCB4 | $^{IF}$ABCB4-(ATP)$_2$ | $^{IF}$ABCB4-PC | $^{IF}$ABCB4-PC-(ATP)$_2$ | $^{CC}$ABCB4-(ATP)$_2$ |
|---|---|---|---|---|---|---|---|
| **Cholesterol** | **Cholesterol** (Lipid 21 residue name: CHL) | **inner** | 54 | 54 | 55 | 56 | 57 |
| | | **outer** | 57 | 57 | 60 | 59 | 59 |
| **POPC** | **1-palmitoyl-2-oleyl-sn-glycero-3-phosphatidylcholine** (Lipid21 sequence: PA-PC-OL) | **inner** | 108 | 108 | 109 | 108 | 108 |
| | | **outer** | 111 | 111 | 109 | 110 | 110 |
| **POPE** | **1-palmitoyl-2-oleyl-sn-glycero-3-phosphatidylcholine** (Lipid21 sequence: PA-PE-OL) | **inner** | 54 | 54 | 54 | 54 | 54 |
| | | **outer** | 55 | 55 | 54 | 54 | 54 |

**Supporting Table S2. Lipid composition (number of lipids) for asymmetric membrane models.**

| Lipids | Full name (Lipid21 residue topology) | Leaflet | apo $^{IF}$ABCB4 | $^{IF}$ABCB4-(ATP)$_2$ | $^{IF}$ABCB4-PC | $^{IF}$ABCB4-PC-(ATP)$_2$ | $^{CC}$ABCB4-(ATP)$_2$ |
|---|---|---|---|---|---|---|---|
| **Cholesterol** | **Cholesterol** (Lipid 21 residue name: CHL) | **inner** | 75 | 75 | 79 | 79 | 82 |
| | | **outer** | 90 | 90 | 87 | 87 | 92 |
| **POPC** | **1-palmitoyl-2-oleyl-sn-glycero-3-phosphatidylcholine** (Lipid21 sequence: PA/PC/OL) | **inner** | 40 | 40 | 41 | 41 | 41 |
| | | **outer** | 96 | 96 | 101 | 101 | 98 |
| **POPE** | **1-palmitoyl-2-oleyl-sn-glycero-3-phosphatidylethanolamine** (Lipid21 sequence: PA/PE/OL) | **inner** | 27 | 27 | 28 | 28 | 28 |
| | | **outer** | 9 | 9 | 10 | 10 | 11 |
| **POPA** | **1-palmitoyl-2-oleyl-sn-glycero-3-phosphatidic acid** (Lipid21 sequence: PA/PH-/OL) | **inner** | 3 | 3 | 3 | 3 | 2 |
| | | **outer** | 0 | 0 | 9 | 9 | 0 |
| **DOPC** | **1,2-dioleyl-sn-glycero-3-phosphatidylcholine** (Lipid21 sequence: OL/PC/OL) | **inner** | 1 | 1 | 1 | 1 | 1 |
| | | **outer** | 3 | 3 | 3 | 3 | 3 |
| **DOPE** | **1,2-dioleyl-sn-glycero-3-phosphatidylethanolamine** (Lipid21 sequence: OL/PE/OL) | **inner** | 6 | 6 | 6 | 6 | 5 |
| | | **outer** | 1 | 1 | 1 | 1 | 1 |
| **POPS** | **1-palmitoyl-2-oleyl-sn-glycero-3-phospho-L-serine** (Lipid21 sequence: PA/PS/OL) | **inner** | 8 | 8 | 8 | 8 | 8 |
| | | **outer** | 0 | 0 | 0 | 0 | 0 |
| **PAPA** | **1-palmitoyl-2-arachidonoyl-sn-glycero-3-phosphatidic acid** (Lipid21 sequence: PA/PH-/AR) | **inner** | 1 | 1 | 1 | 1 | 1 |
| | | **outer** | 0 | 0 | 0 | 0 | 0 |
| **PAPC** | **1-palmitoyl-2-arachidonoyl-sn-glycero-3-phosphatidylcholine** (Lipid21 sequence: PA/PC/AR) | **inner** | 4 | 4 | 4 | 4 | 4 |
| | | **outer** | 8 | 8 | 8 | 8 | 8 |
| **PAPE** | **1-palmitoyl-2-arachidonoyl-sn-glycero-3-phosphatidylethanolamine** (Lipid21 sequence: PA/PE/AR) | **inner** | 17 | 17 | 18 | 18 | 18 |
| | | **outer** | 4 | 4 | 4 | 4 | 4 |

**Supporting Table S2.** *(continued)*

| Lipids | Full name (Lipid21 residue topology) | Leaflet | apo $^{IF}$ABCB4 | $^{IF}$ABCB4-(ATP)$_2$ | $^{IF}$ABCB4-PC | $^{IF}$ABCB4-PC-(ATP)$_2$ | $^{CC}$ABCB4-(ATP)$_2$ |
|---|---|---|---|---|---|---|---|
| **PAPS** | **1-palmitoyl-2-arachidonoyl-sn-glycero-3-phospho-L-serine** (Lipid21 sequence: PA/PS/AR) | **inner** | 14 | 14 | 15 | 15 | 15 |
| | | **outer** | 0 | 0 | 0 | 0 | 0 |
| **PDoPC** | **1-palmitoyl-2-Docosahexaenoyl-sn-glycero-3-phosphatidylcholine** (Lipid21 sequence: PA/PC/DHA) | **inner** | 1 | 1 | 1 | 1 | 1 |
| | | **outer** | 2 | 2 | 2 | 2 | 2 |
| **PDoPE** | **1-palmitoyl-2-docosahexaenoyl-sn-glycero-3-phosphatidylethanolamine** (Lipid21 sequence: PA/PE/DHA) | **inner** | 5 | 5 | 6 | 6 | 5 |
| | | **outer** | 1 | 1 | 1 | 1 | 1 |
| **DAPC** | **1,2-diarachidonoyl-sn-glycero-3-phosphatidylcholine** (Lipid21 sequence: AR/PC/AR) | **inner** | 0 | 0 | 0 | 0 | 0 |
| | | **outer** | 1 | 1 | 1 | 1 | 1 |
| **DAPE** | **1,2-diarachidonoyl-sn-glycero-3-phosphatidylcholine** (Lipid21 sequence: AR/PE/AR) | **inner** | 10 | 10 | 10 | 10 | 9 |
| | | **outer** | 2 | 2 | 2 | 2 | 2 |
| **DDoPE** | **1,2-didocosahexaenoyl-sn-glycero-3-phosphatidylethanolamine** (Lipid21 sequence: DHA/PE/DHA) | **inner** | 3 | 3 | 3 | 3 | 3 |
| | | **outer** | 1 | 1 | 1 | 1 | 0 |
| **DDoPS** | **1,2-didocosahexaenoyl-sn-glycero-3- phospho-L-serine** (Lipid21 sequence: DHA/PS/DHA) | **inner** | 1 | 1 | 1 | 1 | 1 |
| | | **outer** | 0 | 0 | 0 | 0 | 0 |

**Supporting Table S3. System details MD simulations conducted in the present study.**

| Membranes | System setup | apo $^{IF}$ABCB4 | $^{IF}$ABCB4-$(ATP)_2$ | $^{IF}$ABCB4-PC | $^{IF}$ABCB4-PC-$(ATP)_2$ | $^{CC}$ABCB4-$(ATP)_2$ |
|---|---|---|---|---|---|---|
| **Symmetric** | Box Size (x, Å) | 123.06 | 123.06 | 121.55 | 121.55 | 122.68 |
| | Box Size (y, Å) | 123.27 | 123.27 | 126.37 | 126.37 | 124.75 |
| | Box Size (z, Å) | 182.58 | 182.58 | 182.83 | 182.83 | 184.85 |
| | $N_{atoms}$ | 308798 | 249042 | 249905 | 249989 | 252765 |
| | $N_{water}$ | 59840 | 59840 | 60144 | 60144 | 61042 |
| **Asymmetric** | Box Size (x, Å) | 121.92 | 121.92 | 121.64 | 121.64 | 123.41 |
| | Box Size (y, Å) | 118.54 | 118.54 | 1221.68 | 1221.68 | 122.62 |
| | Box Size (z, Å) | 182.58 | 182.58 | 182.83 | 182.83 | 184.855 |
| | $N_{atoms}$ | 292171 | 235662 | 243581 | 243665 | 242090 |
| | $N_{water}$ | 59840 | 56593 | 58586 | 58586 | 58385 |

**Supporting Table S4. MD simulation time (μs) for each replica, conformation and bound state.**

| Membrane | | apo $^{IF}$ABCB4 | $^{IF}$ABCB4-$(ATP)_2$ | $^{IF}$ABCB4-PC | $^{IF}$ABCB4-PC-$(ATP)_2$ | $^{CC}$ABCB4-$(ATP)_2$ |
|---|---|---|---|---|---|---|
| **Symmetric** | **# replicas** | 3 | 3 | 3 | 3 | 3 |
| | **MD Time** | 2 | 2 | 2 | 2 | 2 |
| | **Aggregated MD time** | 6 | 6 | 6 | 6 | 6 |
| **Asymmetric** | **# replicas** | 3 | 3 | 3 | 3 | 3 |
| | **MD Time** | 2 | 2 | 2 | 2 | 2 |
| | **Aggregated MD time** | 6 | 6 | 6 | 6 | 6 |
| **Total Aggregated time** | | 12 | 12 | 12 | 12 | 12 |
| | | 60 μs | | | | |

**Supporting Table S5. ABCB4 topology used in the present manuscript.**

| Domain | Residues |
|---|---|
| TMH1 | 44-89 |
| TMH2 | 108-159 |
| TMH3 | 169-212 |
| TMH4 | 215-261 |
| TMH5 | 271-324 |
| TMH6 | 330-371 |
| TMH7 | 707-739 |
| TMH8 | 747-797 |
| TMH9 | 809-852 |
| TMH10 | 855-901 |
| TMH11 | 912-965 |
| TMH12 | 969-1012 |
| NBD1 | 392-612 |
| NBD2 | 1051-1244 |

**Supporting Table S6. Residues selected for calculations of ABC structural parameters of ABCB4, namely a) EC and IC angles, b) NBD distance and NBD twist.**

(a)

| EC angle | | IC angle | |
|---|---|---|---|
| **Vector1 direction** | **Vector2 direction** | **Vector1 direction** | **Vector2 direction** |
| Chain A 319-321,332-334,733-737,748-752 | Chain A 77-80,109-116,957-960,971-974 | Chain A 52-53,140-158,173-188,358-370,877-900,914-933 | Chain A 236-261,275-294,777-793,812-829,996-1009 |

(b)

| NBD distance | | NBD twist angle | | | |
|---|---|---|---|---|---|
| ***NBD1*** | ***NBD2*** | ***Vector1 origin*** | ***Vector1 direction*** | ***Vector2 origin*** | ***Vector2 direction*** |
| Chain A 393-605 | Chain A 1034-1248 | Chain A 393-476,553-558,583-612 | Chain A 481-549,565-579 | Chain A 1034-1116,1195-1200,1225-1248 | Chain A 1121-1191,1207-1222 |

**Supporting Table S7. ABC structural parameters of resolved ABC proteins used in the present study for ABC conformational space. Conformation, bound- and nucleotide-bound states, PDB IDs are reported as well as extracellular angle (EC, deg), intracellular angle (IC, deg), NBD distance (NBD dist, Å) and NBD twist (deg).**

| Protein | Conformation | Bound-State | Nucleotide | pdb | EC | IC | NBD dist | NBD twist |
|---|---|---|---|---|---|---|---|---|
| MRP1 | IF | LIG | noATP | 5uja | 15.3 | 29.6 | 43.3 | -150.1 |
| ABCC4 | IF | LIG | noATP | 8BWP | 15.0 | 31.6 | 48.1 | -152.7 |
| ABCB8 | IF | APO | 2ATP | 7EHL | 15.3 | 43.1 | 50.9 | -146.4 |
| ABCB1 | IF | LIG | noATP | 6QEX | 12.6 | 34.3 | 38.4 | -144.8 |
| ABCB1 | IF | INH | noATP | 7A6E | 14.4 | 44.2 | 55.5 | -153.0 |
| ABCB8 | OF | APO | 2ADP | 5OCH | 27.9 | 31.1 | 31.6 | -175.9 |
| ABCB7 | IF | APO | AMP-PNP | 7VGF | 12.3 | 33.9 | 54.7 | -157.0 |
| ABCB9 | OF | APO | ATP-ADP | 7V5C | 26.7 | 18.3 | 28.5 | -159.0 |
| ABCB9 | IF | LIG | noATP | 7V5D | 12.6 | 40.3 | 51.5 | -146.1 |
| ABCC4 | IF | LIG | noATP | 8BWR | 14.9 | 31.9 | 48.0 | -155.8 |
| ABCB4 | IF | APO | noATP | 7NIU | 14.9 | 31.1 | 38.3 | -146.4 |
| ABCB11 | OF | APO | 2ATP | 8PMD | 17.6 | 20.7 | 27.7 | -157.9 |
| ABCB4 | IF | LIG | noATP | 7NIV | 13.8 | 34.1 | 37.7 | -144.7 |
| ABCB11 | IF | LIG | noATP | 7E1A | 14.8 | 36.5 | 46.4 | -151.5 |
| MRP1 | IF | INH | noATP | 8F4B | 15.4 | 36.7 | 54.5 | -160.2 |
| ABCB4 | IF | INH | noATP | 7NIW | 13.4 | 36.1 | 40.7 | -148.2 |
| ABCC4 | IF | APO | noATP | 8BJF | 14.8 | 31.8 | 47.6 | -149.1 |
| ABCB1L335C | OF | INH | 2ATP | 7ZK5 | 10.3 | 20.2 | 24.2 | -162.7 |
| bacPCAT1 | IF | APO | noATP | 4RY2 | 15.1 | 27.8 | 33.0 | -143.8 |
| bacMcjD | OF | APO | noATP | 4PL0 | 13.0 | 22.2 | 27.6 | -146.9 |
| bacTmrAB | OF | APO | ATP-ADP | 6RAK | 14.1 | 22.1 | 28.0 | -157.1 |
| bMRP1 | OF | APO | 2ATP | 6BHU | 16.7 | 21.6 | 27.5 | -149.6 |
| bacTmrAB | IF | APO | 2SO4 | 5MKK | 15.2 | 33.1 | 38.9 | -139.2 |
| bacMsbA | OF | APO | noATP | 5TTP | 13.1 | 21.2 | 28.3 | -159.9 |
| bacTmrAB | IF | APO | 2ADP | 6RAF | 15.0 | 29.9 | 39.0 | -139.8 |
| ABCB1 | OF | APO | ATP | 6COV | 15.9 | 20.9 | 28.8 | -157.3 |
| bMRP1 | OF | APO | ATP-ADP | 6UY0 | 17.2 | 21.3 | 28.0 | -149.6 |
| bacTmrAB | OF | APO | 2ATP | 6RAH | 22.4 | 21.2 | 27.9 | -155.0 |
| bacSav | OF | APO | ANP | 2ONJ | 21.1 | 21.7 | 27.4 | -155.0 |
| bacTmrAB | OF | APO | ATP | 6RAI | 13.6 | 22.3 | 28.0 | -156.9 |
| bacTmrAB | IF | APO | noATP | 6RAN | 14.8 | 33.6 | 40.5 | -143.0 |
| bacPCAT1 | OF | APO | ATP-PgS | 4S0F | 15.5 | 20.8 | 27.6 | -151.1 |
| bacTmrAB | IF | APO | ATP-ADP | 6RAG | 15.4 | 32.6 | 39.8 | -141.0 |
| bacTmrAB | OF | APO | ATP-ADP | 6RAJ | 24.0 | 21.7 | 28.0 | -156.1 |
| ABCB4 | OF | APO | 2ATP | 6S7P | 15.7 | 21.0 | 28.1 | -153.7 |
| bacPglK | OF | APO | noATP | 5C73 | 19.5 | 21.9 | 28.0 | -152.1 |
| bacTmrAB | UR | APO | 2ATP | 6RAL | 14.5 | 23.3 | 28.8 | -155.8 |
| bMRP1 | IF | LIG | noATP | 5UJA | 15.3 | 29.6 | 43.3 | -149.7 |
| bacTmrAB | UR | APO | ATP-ADP | 6RAM | 14.7 | 25.4 | 29.5 | -156.2 |

**Supporting Table S8. Distributions of subpopulation identified from InfleCS clustering in (a) symmetric and (b) asymmetric membrane models**

(a)

| Conformations | ABC Parameters | Population IDs | | | | | | | | | |
|---|---|---|---|---|---|---|---|---|---|---|---|
| | | **1** | **2** | **3** | **4** | **5** | **6** | **7** | **8** | **9** | **10** |
| **Apo $^{IF}$ABCB4** | **NBDdist** *vs* **IC** | 47.2% | 3.8% | 20.2% | 4.6% | 15.1% | 9.0% | - | - | - | - |
| | **NBDdist** *vs* **NBD Twist** | 9.7% | 4.7% | 3.7% | 1.4% | 0.7% | 58.6% | 0.1% | 0.4% | 13.1% | 7.5% |
| **$^{IF}$ABCB4-(ATP)$_2$** | **NBDdist** *vs* **IC** | 40.1% | 18.5% | 41.4% | - | - | - | - | - | - | - |
| | **NBDdist** *vs* **NBD Twist** | 22.7% | 64.7% | 3.0% | 1.4% | 8.2% | - | - | - | - | - |
| **$^{IF}$ABCB4-PC** | **NBDdist** *vs* **IC** | 36.5% | 40.3% | 2.9% | 20.3% | - | - | - | - | - | - |
| | **NBDdist** *vs* **NBD Twist** | 9.6% | 15.2% | 9.0% | 35.0% | 1.9% | 29.3% | - | - | - | - |
| **$^{IF}$ABCB4-PC-(ATP)$_2$** | **NBDdist** *vs* **IC** | 37.4% | 14.9% | 2.5% | 14.3% | 4.6% | 11.5% | 9.5% | 5.3% | - | - |
| | **NBDdist** *vs* **NBD Twist** | 23.6% | 7.3% | 36.0% | 16.2% | 13.4% | 3.4% | - | - | - | - |
| **$^{CC}$ABCB4-(ATP)$_2$** | **NBDdist** *vs* **IC** | 30.2% | 26.3% | 43.5% | - | - | - | - | - | - | - |
| | **NBDdist** *vs* **NBD Twist** | 39.8% | 60.2% | - | - | - | - | - | - | - | - |

(b)

| **Conformations** | **ABC Parameters** | **Population IDs** | | | | | | | | | | |
|---|---|---|---|---|---|---|---|---|---|---|---|---|
| | | **1** | **2** | **3** | **4** | **5** | **6** | **7** | **8** | **9** | **10** | **11** |
| **Apo $^{IF}$ABCB4** | **NBDdist** *vs* **IC** | 6.2% | 0.5% | 6.8% | 15.0% | 7.5% | 18.6% | 25.5% | 6.7% | 2.7% | 5.3% | 5.2% |
| | **NBDdist** *vs* **NBD Twist** | 7.1% | 13.7% | 2.5% | 6.4% | 1.4% | 35.4% | 1.1% | 13.3% | 0.1% | 19.0% | - |
| **$^{IF}$ABCB4-(ATP)$_2$** | **NBDdist** *vs* **IC** | 35.8% | 28.8% | 10.3% | 13.2% | 8.5% | 3.5% | - | - | - | - | - |
| | **NBDdist** *vs* **NBD Twist** | 3.1% | 21.6% | 7.6% | 32.7% | 35.0% | - | - | - | - | - | - |
| **$^{IF}$ABCB4-PC** | **NBDdist** *vs* **IC** | 58.0% | 0.8% | 2.8% | 38.3% | - | - | - | - | - | - | - |
| | **NBDdist** *vs* **NBD Twist** | 3.0% | 9.6% | 6.5% | 7.9% | 11.9% | 4.4% | 37.3% | 17.8% | 1.6% | - | - |
| **$^{IF}$ABCB4-PC-(ATP)$_2$** | **NBDdist** *vs* **IC** | 11.5% | 4.2% | 6.6% | 9.8% | 9.7% | 3.6% | 14.7% | 4.2% | 35.6% | - | - |
| | **NBDdist** *vs* **NBD Twist** | 3.2% | 27.9% | 27.1% | 5.7% | 36.1% | - | - | - | - | - | - |
| **$^{CC}$ABCB4-(ATP)$_2$** | **NBDdist** *vs* **IC** | 55.9% | 44.1% | - | - | - | - | - | - | - | - | - |
| | **NBDdist** *vs* **NBD Twist** | 32.1% | 23.5% | 44.4% | - | - | - | - | - | - | - | - |

**Supporting Table S9. Identified sequence-based CRAC and CARC motifs in ABCB4 sequence.**

| Motif | Name | Location (residues) |
|---|---|---|
| CRAC motif | CRAC1 | 44-54 |
| | CRAC2 | 246-251 |
| | CRAC3 | 276-281 |
| | CRAC4 | 982-995 |
| CARC motif | CARC1 | 54-59 |
| | CARC2 | 115-123 |
| | CARC3 | 147-158 |
| | CARC4 | 191-199 |
| | CARC5 | 236-255 |
| | CARC6 | 274-283 |
| | CARC7 | 767-778 |
| | CARC8 | 812-821 |
| | CARC9 | 848-858 |
| | CARC10 | 975-993 |

**Supporting Table S10. Calculated protein-lipid H-bond fractions over MD simulations.** H-bond in gray-shaded cells are H-bond either appearing or strenghtened during MD production as compared with protein equilibration stage (i.e., first 20 ns of production runs). H-bond in red-shaded cells are H-bonds present during the production equilibration (i.e., first 20 ns of production runs) but disappearing along MD simulations. The so-called H-bond fractions were calculated for the last μs of MD production run.

| | | Asymmetric | | | | | Symmetric | | | | |
|---|---|---|---|---|---|---|---|---|---|---|---|
| | | **Apo $^{IF}$ABCB4** | **$^{IF}$ABCB4-(ATP)$_2$** | **$^{IF}$ABCB4-PC** | **$^{IF}$ABCB4-PC-(ATP)$_2$** | **$^{CC}$ABCB4-(ATP)$_2$** | **Apo $^{IF}$ABCB4** | **$^{IF}$ABCB4-(ATP)$_2$** | **$^{IF}$ABCB4-PC** | **$^{IF}$ABCB4-PC-(ATP)$_2$** | **$^{CC}$ABCB4-(ATP)$_2$** |
| **Thr44** | | **0.71** | 0.70 | 0.52 | 1.08 | **1.39** | **0.72** | 1.00 | **0.83** | **1.01** | **0.89** |
| **Leu45** | | 0.37 | 0.00 | 0.00 | 0.00 | 0.00 | 0.00 | 0.00 | 0.00 | 0.00 | **0.41** |
| **Arg47** | | 0.00 | 0.00 | 0.00 | 0.00 | 0.59 | 0.00 | 0.00 | 0.00 | 0.00 | 0.00 |
| **Ser49** | | 0.00 | 0.00 | 0.00 | 0.00 | 0.00 | 0.00 | 0.00 | 0.00 | 0.00 | **0.59** |
| **Trp51** | **TMH1** | 0.00 | 0.00 | 0.00 | 0.00 | 0.00 | 0.00 | 0.00 | 0.00 | **0.40** | 0.00 |
| **Gln52** | **TMH1** | **0.40** | 0.51 | 0.45 | **0.87** | 0.00 | 0.35 | **0.52** | **0.41** | 0.40 | **0.47** |
| **Lys54** | **TMH1** | 0.64 | **1.18** | **1.06** | **0.89** | **0.84** | 0.55 | **1.06** | **0.86** | **1.22** | 1.01 |
| **Arg115** | **TMH2** | **0.76** | 1.21 | **1.15** | **1.27** | **1.03** | **0.96** | 1.25 | **1.17** | **1.22** | **0.71** |
| **Tyr116** | **TMH2** | 0.00 | 0.00 | 0.00 | 0.00 | 0.00 | 0.00 | **0.36** | 0.00 | 0.00 | 0.00 |
| **Tyr119** | **TMH2** | 0.00 | 0.00 | 0.00 | 0.00 | **0.61** | 0.00 | **0.49** | 0.00 | 0.00 | 0.00 |
| **Arg144** | **TMH2** | 0.56 | **1.15** | 1.07 | **1.35** | **0.54** | **1.03** | **0.67** | **1.02** | 0.77 | **1.15** |
| **Arg147** | **TMH2** | **0.55** | **0.60** | 0.52 | **0.44** | 0.00 | 0.00 | 0.00 | 0.00 | 0.00 | 0.00 |
| **Arg212** | **TMH3** | 0.00 | 0.00 | **0.35** | 0.00 | 0.00 | 0.00 | 0.00 | **0.41** | 0.00 | 0.00 |
| **Lys215** | **TMH4** | **0.36** | 0.64 | 0.67 | 0.59 | 0.70 | **0.46** | **1.04** | 0.77 | 0.57 | 0.63 |
| **Lys236** | **TMH4** | 0.71 | 0.97 | **0.40** | **0.46** | **1.15** | 0.78 | **1.13** | **0.41** | **0.57** | **1.05** |
| **Ser239** | **TMH4** | 0.00 | 0.00 | 0.00 | 0.00 | 0.00 | 0.00 | **0.54** | 0.00 | 0.00 | 0.00 |
| **Asp243** | **TMH4** | 0.00 | **0.68** | 0.00 | 0.00 | 0.00 | 0.00 | **0.56** | 0.00 | 0.00 | 0.00 |
| **Glu245** | **TMH4** | 0.00 | 0.00 | 0.00 | **0.38** | 0.00 | 0.00 | 0.00 | 0.00 | 0.00 | 0.00 |
| **Glu288** | **TMH5** | 0.00 | **0.36** | 0.00 | **0.45** | 0.00 | 0.00 | 0.00 | 0.00 | 0.00 | 0.00 |
| **Lys292** | **TMH5** | 0.39 | **0.76** | **0.97** | **0.79** | **0.63** | **0.45** | **0.55** | 0.81 | **0.97** | **0.74** |
| **Lys293** | **TMH5** | 0.00 | 0.00 | 0.47 | 0.45 | 0.00 | 0.00 | 0.00 | 0.38 | 0.42 | 0.00 |
| **Arg361** | **TMH6** | **2.00** | 2.33 | **2.46** | **2.47** | **1.42** | **0.77** | **2.39** | **2.72** | **2.44** | **2.91** |
| **Tyr365** | **TMH6** | 0.00 | **0.66** | **0.54** | **0.61** | 0.00 | 0.00 | **0.55** | **0.35** | **0.53** | 0.00 |
| **Val692** | | 0.00 | 0.00 | 0.00 | **0.38** | 0.00 | 0.00 | 0.00 | 0.00 | 0.00 | **0.38** |
| **Ser696** | | **0.52** | **0.55** | **0.68** | 0.00 | 0.55 | **0.42** | **0.82** | **0.56** | 0.46 | **1.00** |
| **Lys702** | | 0.48 | **0.75** | 0.00 | 0.48 | **0.93** | **0.37** | **0.65** | **0.86** | **0.64** | 0.74 |
| **Asn704** | | 0.00 | 0.00 | **0.46** | 0.00 | **0.35** | | | | | |
| **Lys705** | | 0.00 | 0.77 | 0.64 | 0.72 | **0.43** | 0.00 | **0.51** | **0.60** | 0.00 | **0.65** |

| | | | | | | | | | | | |
|---|---|---|---|---|---|---|---|---|---|---|---|
| **Tyr710** | **TMH7** | 0.00 | 0.00 | 0.00 | **0.52** | 0.00 | 0.00 | **0.40** | 0.00 | 0.41 | 0.00 |
| **Gln749** | **TMH8** | 0.00 | 0.00 | 0.00 | 0.00 | 0.00 | 0.00 | **0.40** | 0.00 | 0.00 | **0.37** |
| **Lys750** | **TMH8** | 0.00 | 0.40 | 0.00 | 0.00 | 0.00 | 0.00 | **0.51** | 0.00 | 0.00 | 0.00 |
| **Asn752** | **TMH8** | 0.00 | 0.00 | 0.00 | 0.00 | 0.00 | 0.00 | 0.52 | 0.00 | 0.00 | **0.41** |
| **Lys778** | **TMH8** | **0.59** | **1.11** | **1.00** | **0.92** | **0.44** | 0.49 | **0.50** | **0.83** | 0.80 | **0.49** |
| **Arg785** | **TMH8** | 0.00 | **0.54** | 0.00 | **0.65** | 0.00 | 0.00 | 0.00 | 0.00 | 0.00 | 0.00 |
| **Arg831** | **TMH9** | 0.43 | 0.00 | **1.85** | **1.67** | 0.00 | 0.00 | 0.38 | **0.42** | **0.87** | 0.00 |
| **Tyr852** | **TMH9** | 0.00 | 0.00 | **0.35** | 0.00 | 0.00 | 0.00 | 0.00 | 0.00 | 0.00 | 0.00 |
| **Glu874** | **TMH10** | 0.00 | 0.00 | 0.00 | 0.00 | 0.00 | 0.00 | 0.00 | 0.00 | 0.00 | **0.37** |
| **Lys876** | **TMH10** | **0.86** | 1.24 | 0.00 | 0.00 | **1.08** | **0.91** | **1.38** | **0.54** | 0.00 | 0.89 |
| **Asn881** | **TMH10** | 0.00 | 0.00 | 0.00 | **0.51** | 0.00 | 0.00 | 0.00 | **0.88** | **0.51** | 0.00 |
| **Lys883** | **TMH10** | 0.00 | 0.00 | **0.54** | 0.00 | 0.00 | 0.00 | 0.00 | 0.00 | 0.00 | 0.00 |
| **Arg884** | **TMH10** | 0.36 | **1.83** | **1.52** | 1.19 | 1.05 | 0.52 | **1.35** | **1.41** | **1.71** | **1.16** |
| **Tyr924** | **TMH11** | 0.00 | 0.00 | 0.00 | **0.38** | 0.00 | 0.00 | **0.37** | 0.00 | **0.61** | 0.00 |
| **Tyr927** | **TMH11** | 0.00 | 0.00 | 0.56 | 0.43 | 0.00 | 0.00 | **0.58** | **0.49** | **0.74** | **0.64** |
| **Arg928** | **TMH11** | 0.55 | 0.00 | **2.68** | **2.11** | 1.61 | 1.06 | **2.42** | **2.71** | **2.32** | **1.70** |
| **Asn929** | **TMH11** | 0.00 | 0.00 | 0.00 | 0.00 | 0.00 | 0.00 | 0.00 | 0.00 | **0.38** | 0.00 |
| **Ser930** | **TMH11** | **0.74** | **0.51** | 0.00 | 0.00 | 0.00 | 0.00 | 0.00 | 0.00 | 0.00 | 0.00 |
| **Gln932** | **TMH11** | 0.00 | 0.42 | **0.79** | **0.62** | 0.47 | 0.37 | **0.99** | **0.86** | **0.87** | **0.85** |
| **Lys933** | **TMH11** | 0.45 | 0.00 | 0.00 | 0.00 | 0.00 | 0.00 | 0.00 | 0.00 | 0.00 | 0.00 |
| **Hie935** | **TMH11** | 0.00 | 0.00 | 0.00 | 0.00 | 0.00 | 0.00 | 0.00 | 0.00 | 0.00 | **0.37** |
| **Arg957** | **TMH11** | 1.23 | 1.33 | 1.31 | **1.17** | **1.23** | 1.03 | **1.61** | **1.20** | 0.96 | **1.36** |
| **Tyr961** | **TMH11** | 0.00 | 0.00 | 0.37 | **0.36** | **0.38** | 0.00 | 0.37 | **0.36** | 0.37 | **0.36** |
| **Arg969** | **TMH12** | **0.37** | **0.73** | **0.51** | 0.39 | **0.73** | 0.00 | **0.51** | **0.49** | **0.50** | **0.75** |
| **Arg971** | **TMH12** | 0.00 | 0.00 | 0.00 | 0.00 | **0.44** | 0.00 | **0.54** | 0.00 | 0.00 | **0.86** |
| **Hid989** | **TMH12** | 0.00 | 0.00 | 0.00 | 0.35 | 0.00 | 0.00 | 0.00 | 0.00 | 0.00 | 0.00 |
| **Tyr997** | **TMH12** | 0.57 | **0.74** | **0.88** | 0.44 | 0.00 | 0.00 | **0.78** | **0.52** | 0.38 | 0.00 |
| **Lys1001** | **TMH12** | 0.53 | **0.97** | **1.11** | **0.75** | 0.84 | **0.64** | **0.85** | **1.00** | **0.57** | **0.69** |

**Supporting Table S11. Calculated helicities of transmembrane domains over MD simulations for each conformation and bound state of ABCB4.**

| Conformation | Membrane | TMH1 | TMH2 | TMH3 | **TMH4** | TMH5 | TMH6 |
|---|---|---|---|---|---|---|---|
| apo $^{IF}$*h*ABCB4 | Symmetric | 0.90 ± 0.24 | 0.94 ± 0.23 | 0.86 ± 0.29 | 0.89 ± 0.23 | 0.94 ± 0.20 | 0.92 ± 0.22 |
| | Asymmetric | 0.90 ± 0.22 | 0.94 ± 0.23 | 0.85 ± 0.29 | 0.94 ± 0.21 | 0.96 ± 0.19 | 0.92 ± 0.23 |
| $^{IF}$*h*ABCB4-(ATP)$_2$ | Symmetric | 0.90 ± 0.22 | 0.93 ± 0.23 | 0.82 ± 0.29 | 0.86 ± 0.25 | 0.95 ± 0.19 | 0.89 ± 0.23 |
| | Asymmetric | 0.89 ± 0.24 | 0.93 ± 0.23 | 0.86 ± 0.30 | 0.93 ± 0.22 | 0.95 ± 0.19 | 0.90 ± 0.24 |
| $^{IF}$*h*ABCB4-PC | Symmetric | 0.92 ± 0.21 | 0.91 ± 0.27 | 0.87 ± 0.28 | **0.80 ± 0.32** | 0.95 ± 0.20 | 0.92 ± 0.23 |
| | Asymmetric | 0.92 ± 0.22 | 0.93 ± 0.23 | 0.86 ± 0.28 | **0.81 ± 0.32** | 0.94 ± 0.20 | 0.92 ± 0.23 |
| $^{IF}$*h*ABCB4-PC-(ATP)$_2$ | Symmetric | 0.93 ± 0.21 | 0.93 ± 0.24 | 0.87 ± 0.28 | **0.79 ± 0.33** | 0.95 ± 0.19 | 0.91 ± 0.26 |
| | Asymmetric | 0.93 ± 0.21 | 0.93 ± 0.24 | 0.87 ± 0.28 | **0.78 ± 0.36** | 0.94 ± 0.20 | 0.92 ± 0.23 |
| $^{CC}$*h*ABCB4-(ATP)$_2$ | Symmetric | 0.82 ± 0.32 | 0.93 ± 0.23 | 0.86 ± 0.27 | 0.93 ± 0.22 | 0.93 ± 0.22 | 0.90 ± 0.22 |
| | Asymmetric | 0.88 ± 0.21 | 0.94 ± 0.22 | 0.83 ± 0.28 | 0.93 ± 0.22 | 0.95 ± 0.20 | 0.90 ± 0.25 |

| Conformation | Membrane | TMH7 | TMH8 | TMH9 | **TMH10** | TMH11 | TMH12 |
|---|---|---|---|---|---|---|---|
| apo $^{IF}$*h*ABCB4 | Symmetric | 0.90 ± 0.24 | 0.91 ± 0.27 | 0.89 ± 0.24 | 0.90 ± 0.22 | 0.95 ± 0.20 | 0.83 ± 0.26 |
| | Asymmetric | 0.90 ± 0.26 | 0.86 ± 0.30 | 0.87 ± 0.25 | 0.85 ± 0.25 | 0.78 ± 0.34 | 0.90 ± 0.24 |
| $^{IF}$*h*ABCB4-(ATP)$_2$ | Symmetric | 0.89 ± 0.24 | 0.88 ± 0.29 | 0.88 ± 0.23 | 0.88 ± 0.22 | 0.94 ± 0.21 | 0.81 ± 0.27 |
| | Asymmetric | 0.91 ± 0.25 | 0.89 ± 0.28 | 0.87 ± 0.26 | 0.83 ± 0.25 | 0.85 ± 0.28 | 0.89 ± 0.23 |
| $^{IF}$*h*ABCB4-PC | Symmetric | 0.87 ± 0.27 | 0.90 ± 0.28 | 0.90 ± 0.24 | **0.72 ± 0.37** | 0.95 ± 0.20 | 0.79 ± 0.33 |
| | Asymmetric | 0.86 ± 0.27 | 0.90 ± 0.29 | 0.89 ± 0.26 | **0.79 ± 0.28** | 0.93 ± 0.21 | 0.84 ± 0.32 |
| $^{IF}$*h*ABCB4-PC-(ATP)$_2$ | Symmetric | 0.87 ± 0.27 | 0.90 ± 0.28 | 0.89 ± 0.24 | **0.71 ± 0.39** | 0.95 ± 0.20 | 0.81 ± 0.31 |
| | Asymmetric | 0.87 ± 0.26 | 0.90 ± 0.28 | 0.86 ± 0.27 | **0.69 ± 0.35** | 0.94 ± 0.20 | 0.86 ± 0.32 |
| $^{CC}$*h*ABCB4-(ATP)$_2$ | Symmetric | 0.86 ± 0.27 | 0.90 ± 0.28 | 0.89 ± 0.27 | 0.86 ± 0.22 | 0.93 ± 0.21 | 0.85 ± 0.25 |
| | Asymmetric | 0.85 ± 0.27 | 0.90 ± 0.28 | 0.85 ± 0.29 | 0.85 ± 0.24 | 0.94 ± 0.20 | 0.86 ± 0.25 |

**Supporting Table S12. Calculated atomic contact fractions between ABCB4 residues and PC substrate for $^{IF}$ABCB4-PC and $^{IF}$ABCB4-PC-(ATP)$_2$.**

| Residues | Symmetric | | Asymmetric | |
|---|---|---|---|---|
| | $^{IF}$ABCB4-PC | $^{IF}$ABCB4-PC-(ATP)$_2$ | $^{IF}$ABCB4-PC | $^{IF}$ABCB4-PC-(ATP)$_2$ |
| **Trp234** | 12.79 | 26.02 | 9.23 | 29.42 |
| **Phe305** | 32.32 | 17.84 | 17.84 | 4.61 |
| **Phe345** | 10.11 | 22.88 | 30.64 | 26.52 |
| **Gln725** | 26.04 | 13.14 | 22.38 | 22.15 |
| **His989** | 12.24 | 18.98 | 25.63 | 19.63 |

**Supporting Table S13. Calculated H-bond fractions between identified binding pocket residues and PC substrate for $^{IF}$ABCB4-PC and $^{IF}$ABCB4-PC-(ATP)$_2$.**

| Binding pocket Residues | Symmetric | | Asymmetric | |
|---|---|---|---|---|
| | **$^{IF}$ABCB4-PC** | **$^{IF}$ABCB4-PC-(ATP)$_2$** | **$^{IF}$ABCB4-PC** | **$^{IF}$ABCB4-PC-(ATP)$_2$** |
| **Trp234** | 0.00 | 0.00 | 0.00 | 0.00 |
| **Phe305** | 0.00 | 0.00 | 0.00 | 0.00 |
| **Phe345** | 0.00 | 0.00 | 0.00 | 0.00 |
| **Gln725** | 0.33 | 0.27 | 0.07 | 0.15 |
| **His989** | 0.06 | 0.26 | 0.32 | 0.35 |

### Supporting Figures

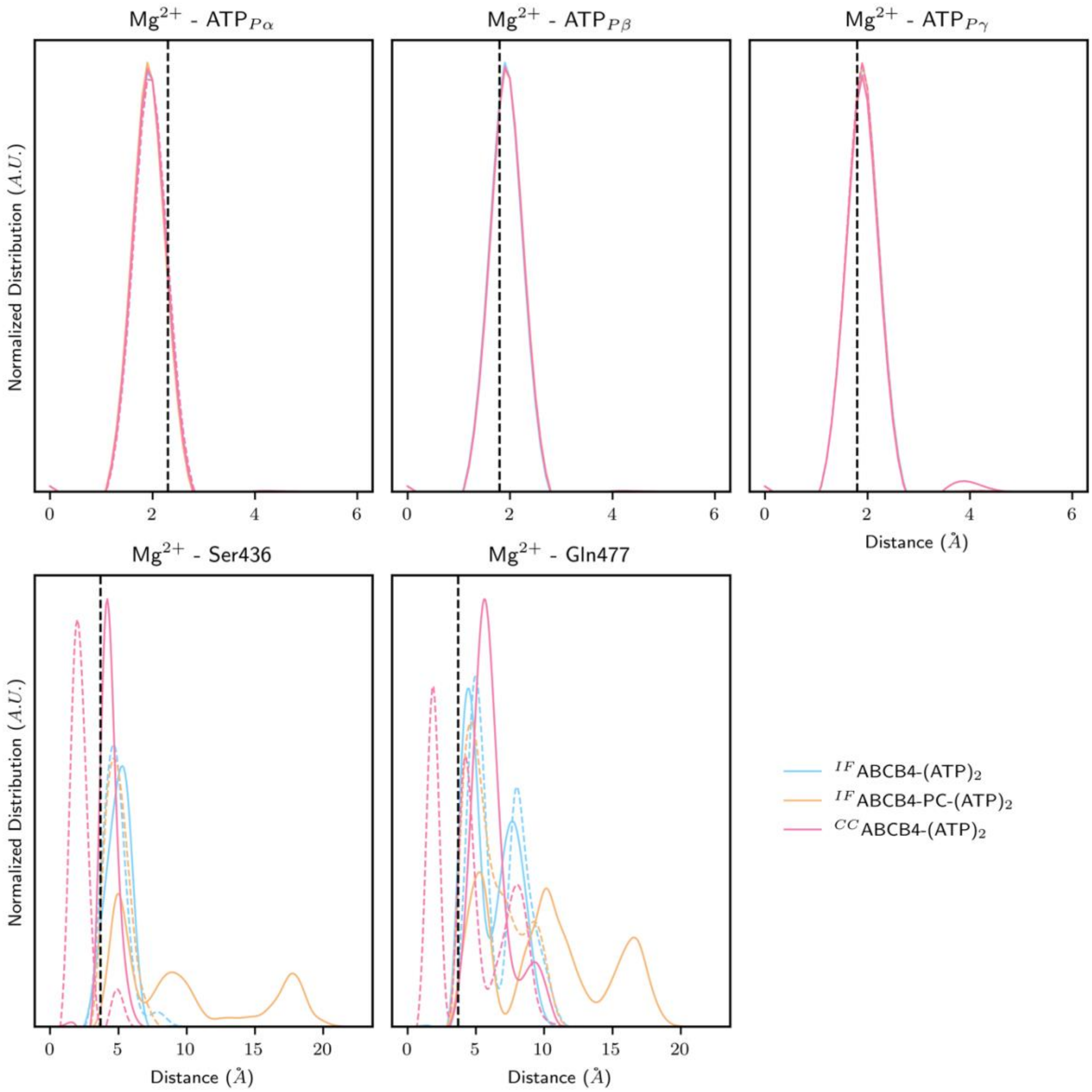

**Supporting Figure S1. Distributions of restrant distances between Mg++ and NBS residues or ATP in NBS1. The distance restraints used in during the equilibration are depicted as vertical black dashed lines.**

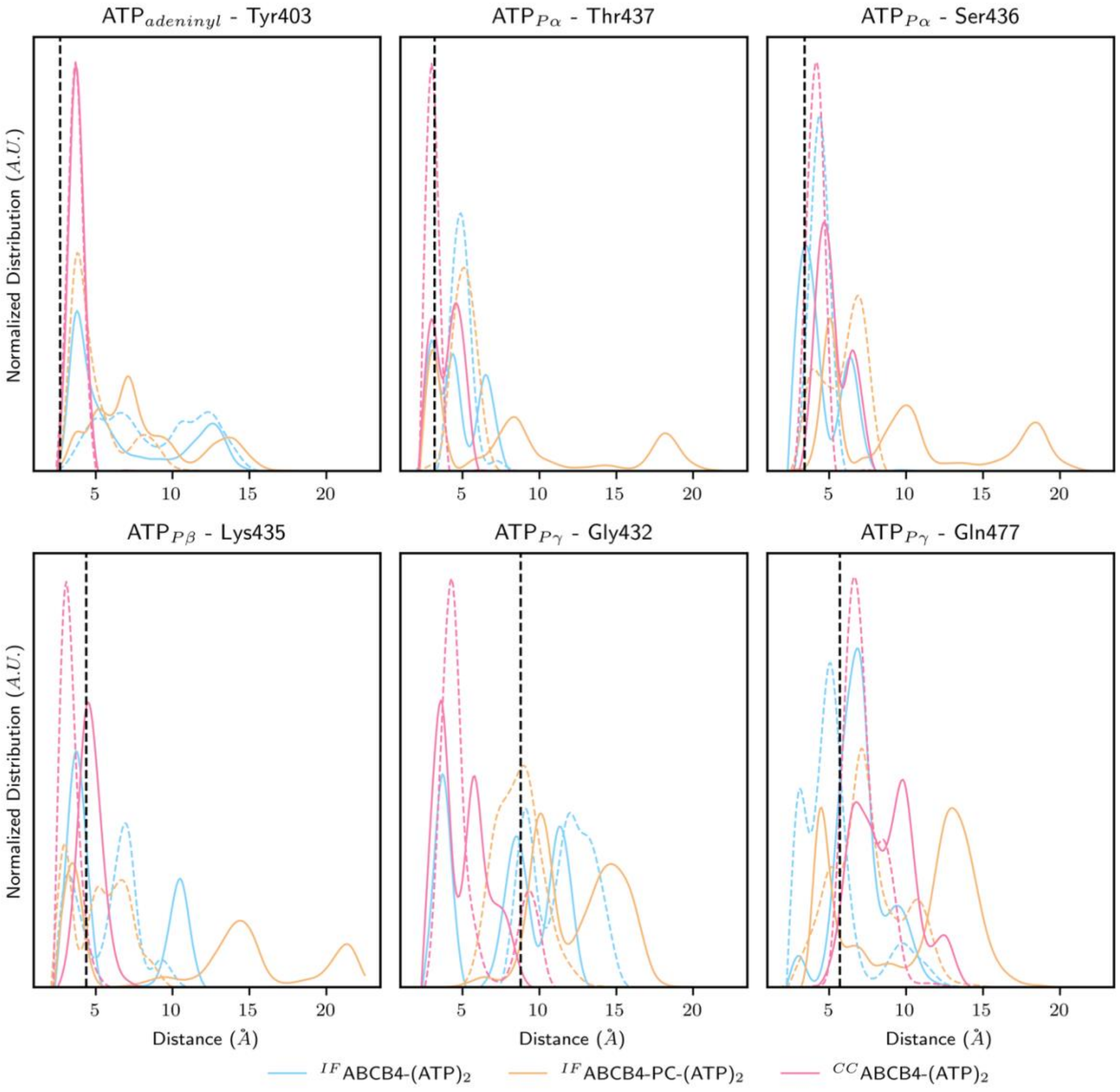

**Supporting Figure S2. Distributions of restrant distances between ATP and NBS residues in NBS1. The distance restraints used in during the equilibration are depicted as vertical black dashed lines.**

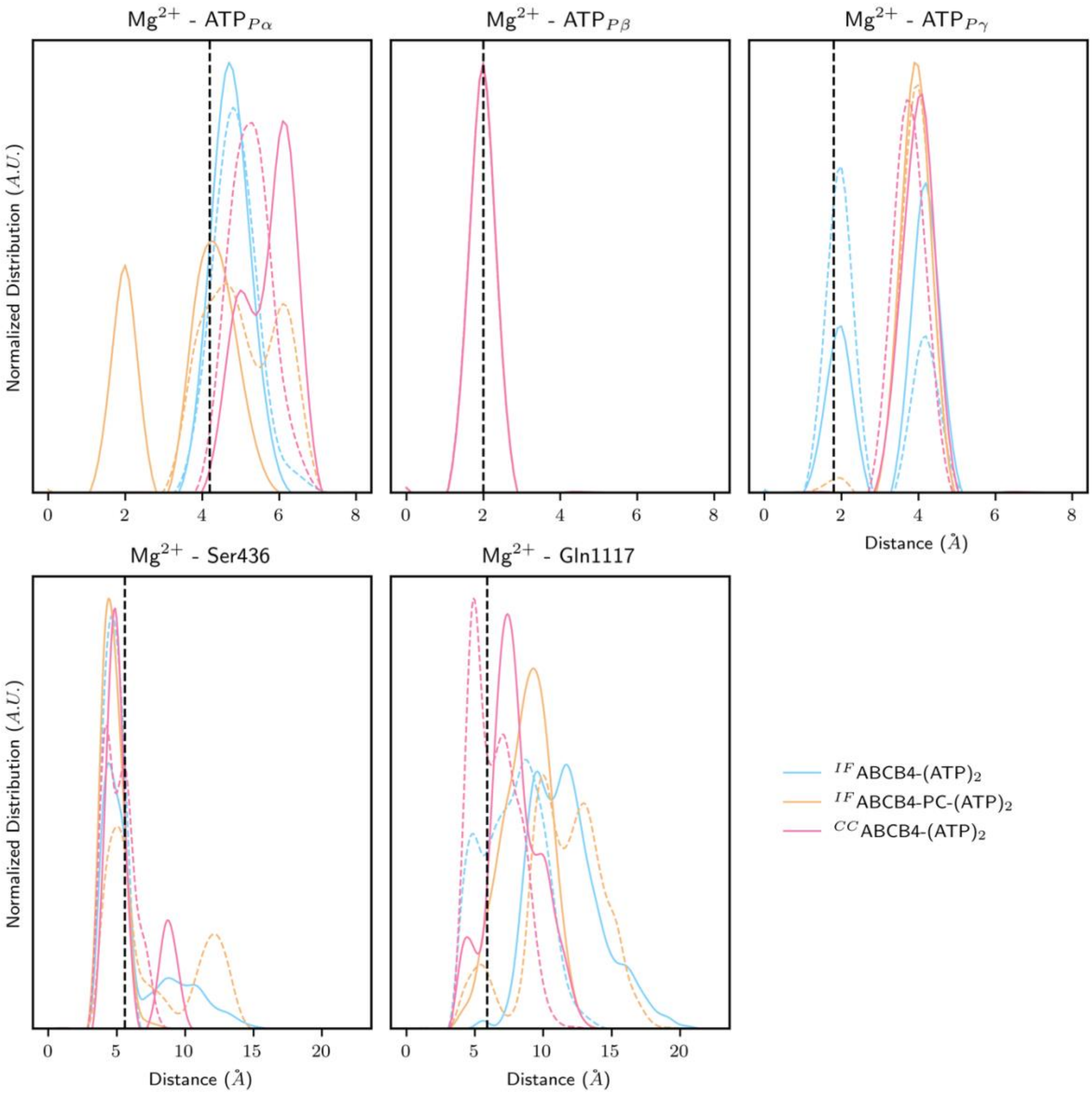

**Supporting Figure S3. Distributions of restrant distances between Mg++ and NBS residues or ATP in NBS2. The distance restraints used in during the equilibration are depicted as vertical black dashed lines.**

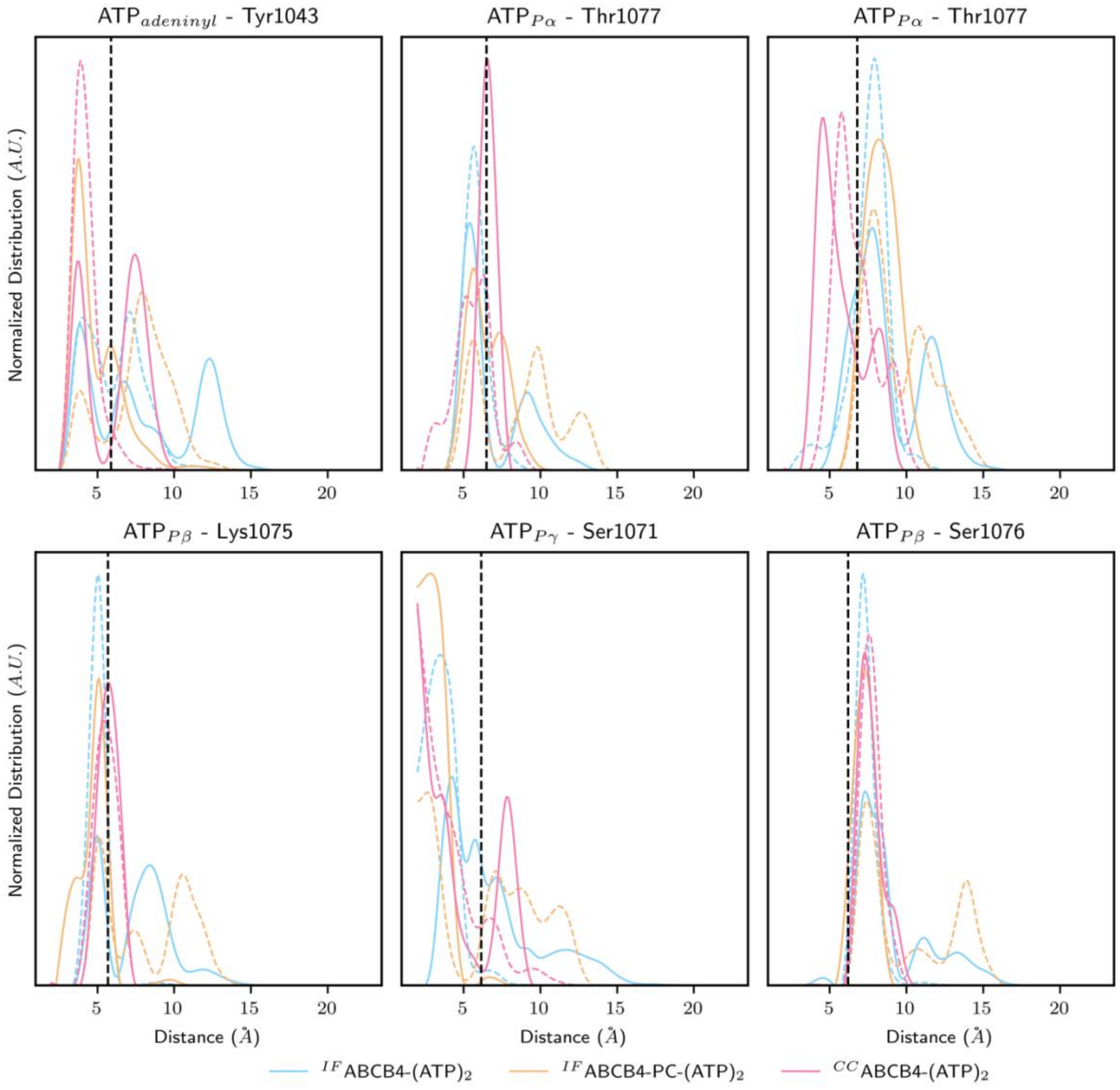

**Supporting Figure S4. Distributions of restrant distances between ATP and NBS residues in NBS2. The distance restraints used in during the equilibration are depicted as vertical black dashed lines.**

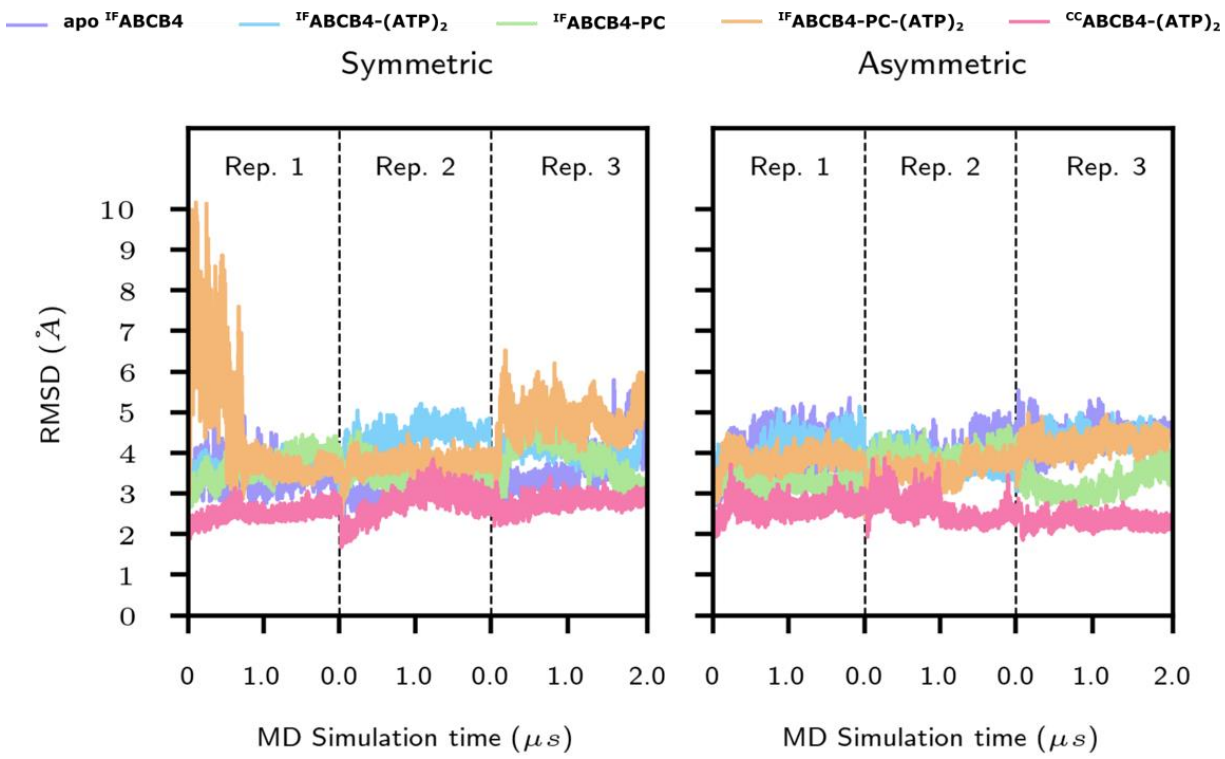

**Supporting Figure S5. Root-mean squared deviation (RMSD, Å) over MD simulation time for each system in symmetric (left) and asymmetric membrane models. RMSDs were calculated over the whole protein using the initial minimized structure as reference.**

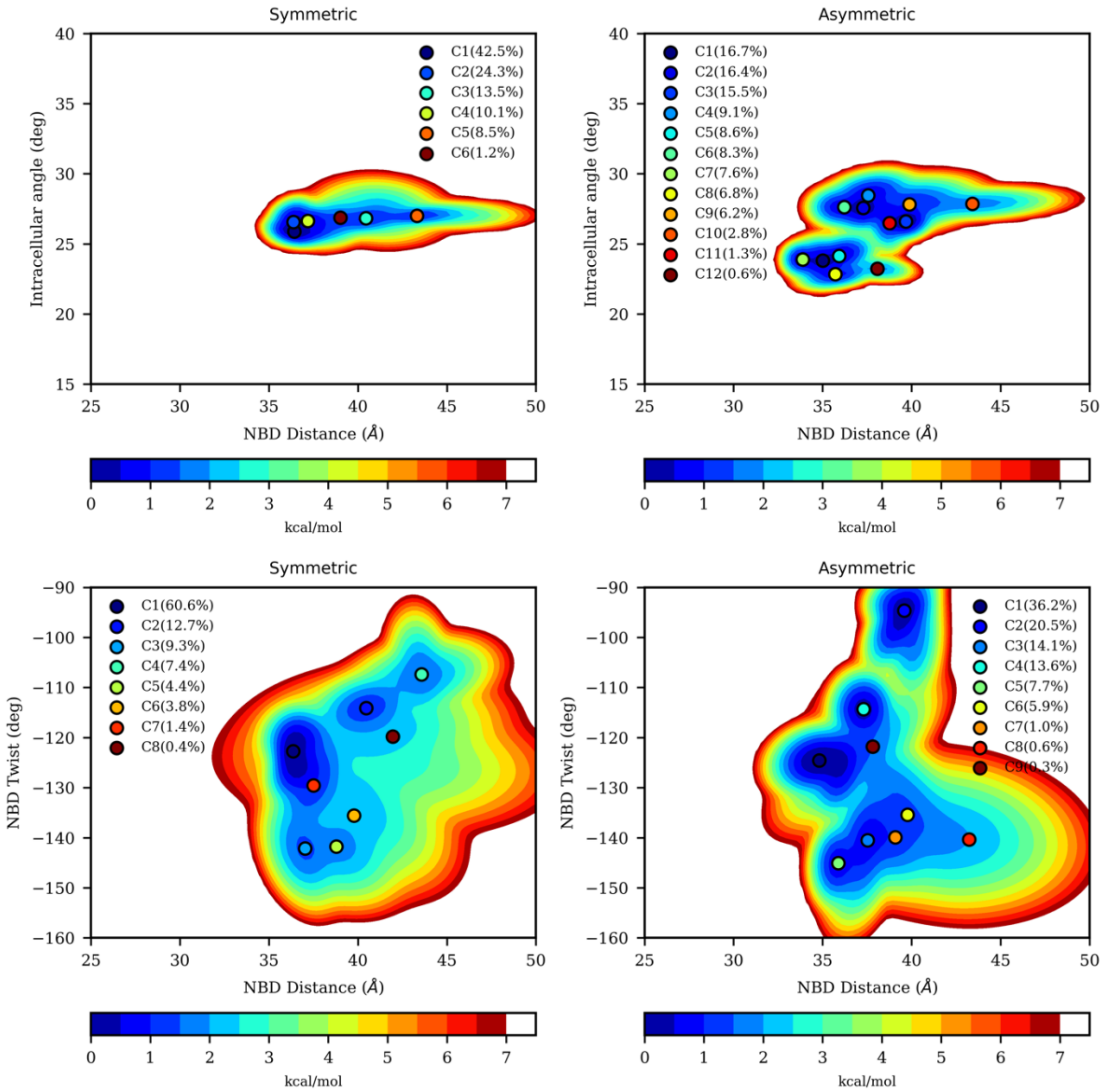

**Supporting Figure S6. Local free energy landscapes calculated for apo IFABCB4 according IC angle vs NBD distance (top) or NBD twist vs NBD distance (bottom) in symmetric (left) and asymmetric (right) membrane model from MD simulations.**

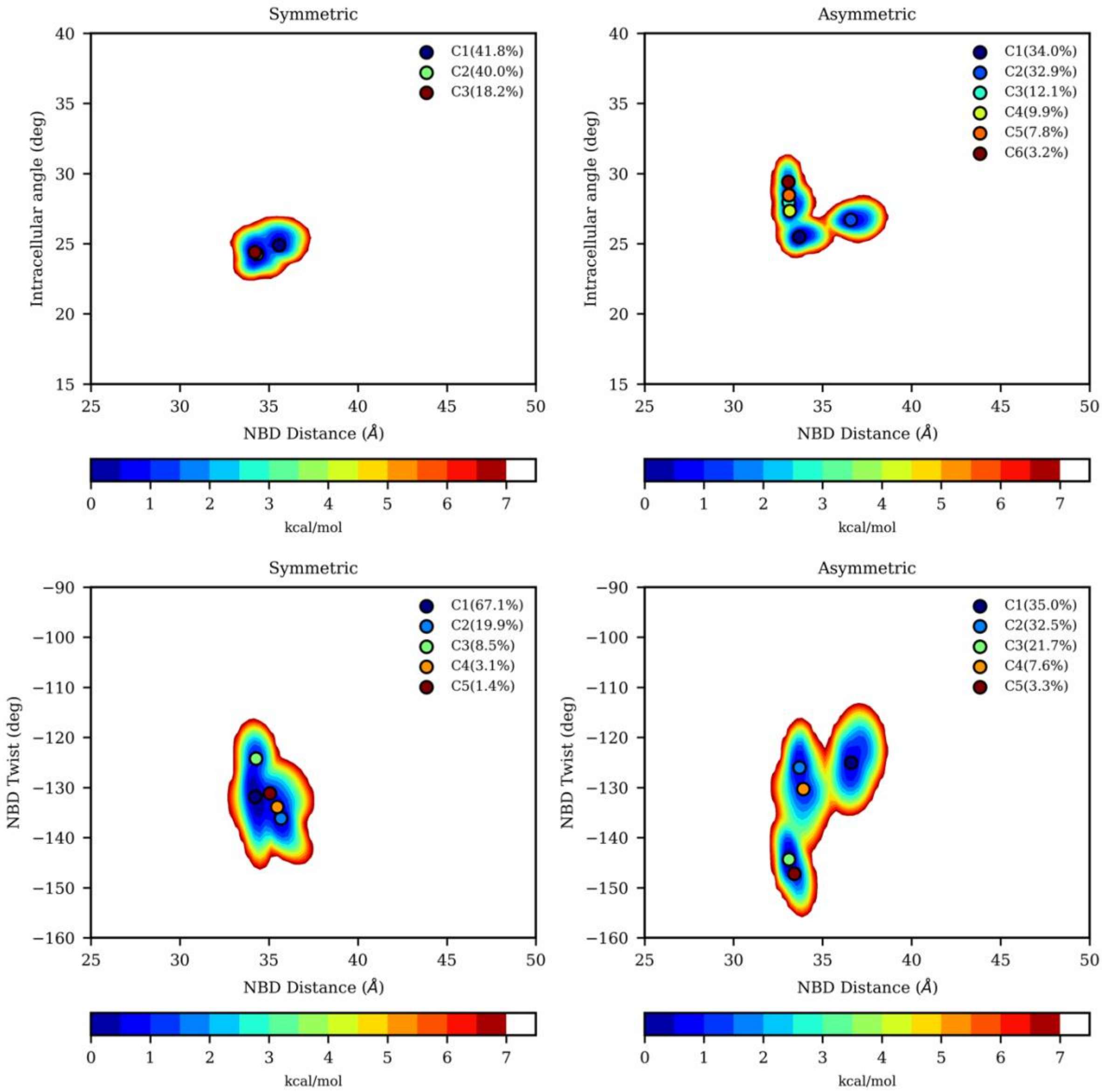

**Supporting Figure S7. Local free energy landscapes calculated for $^{IF}$ABCB4-$(ATP)_2$ according IC angle vs NBD distance (top) or NBD twist vs NBD distance (bottom) in symmetric (left) and asymmetric (right) membrane model from MD simulations.**

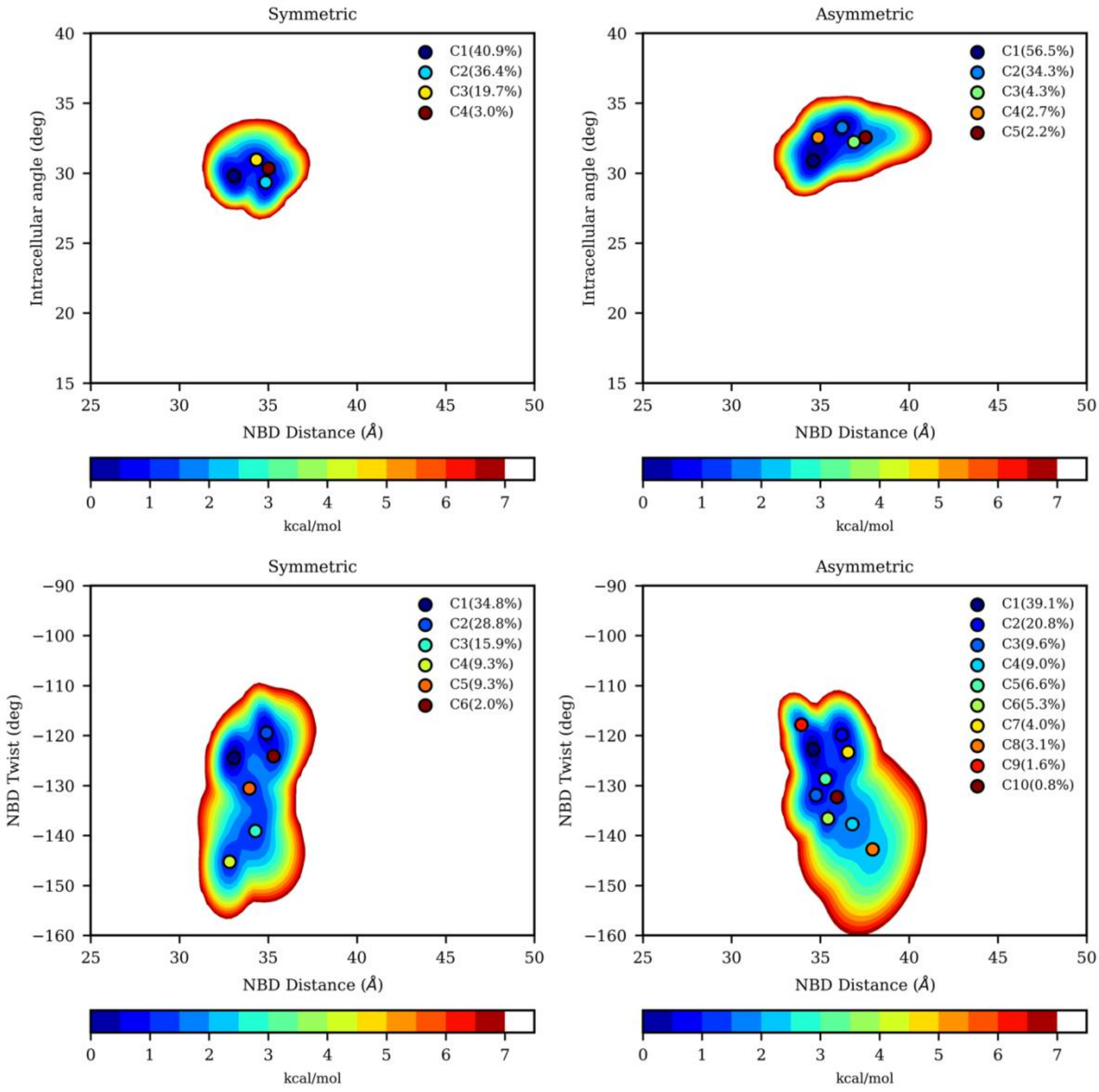

**Supporting Figure S8. Local free energy landscapes calculated for $^{IF}$ABCB4-*PC* according IC angle vs NBD distance (top) or NBD twist vs NBD distance (bottom) in symmetric (left) and asymmetric (right) membrane model from MD simulations.**

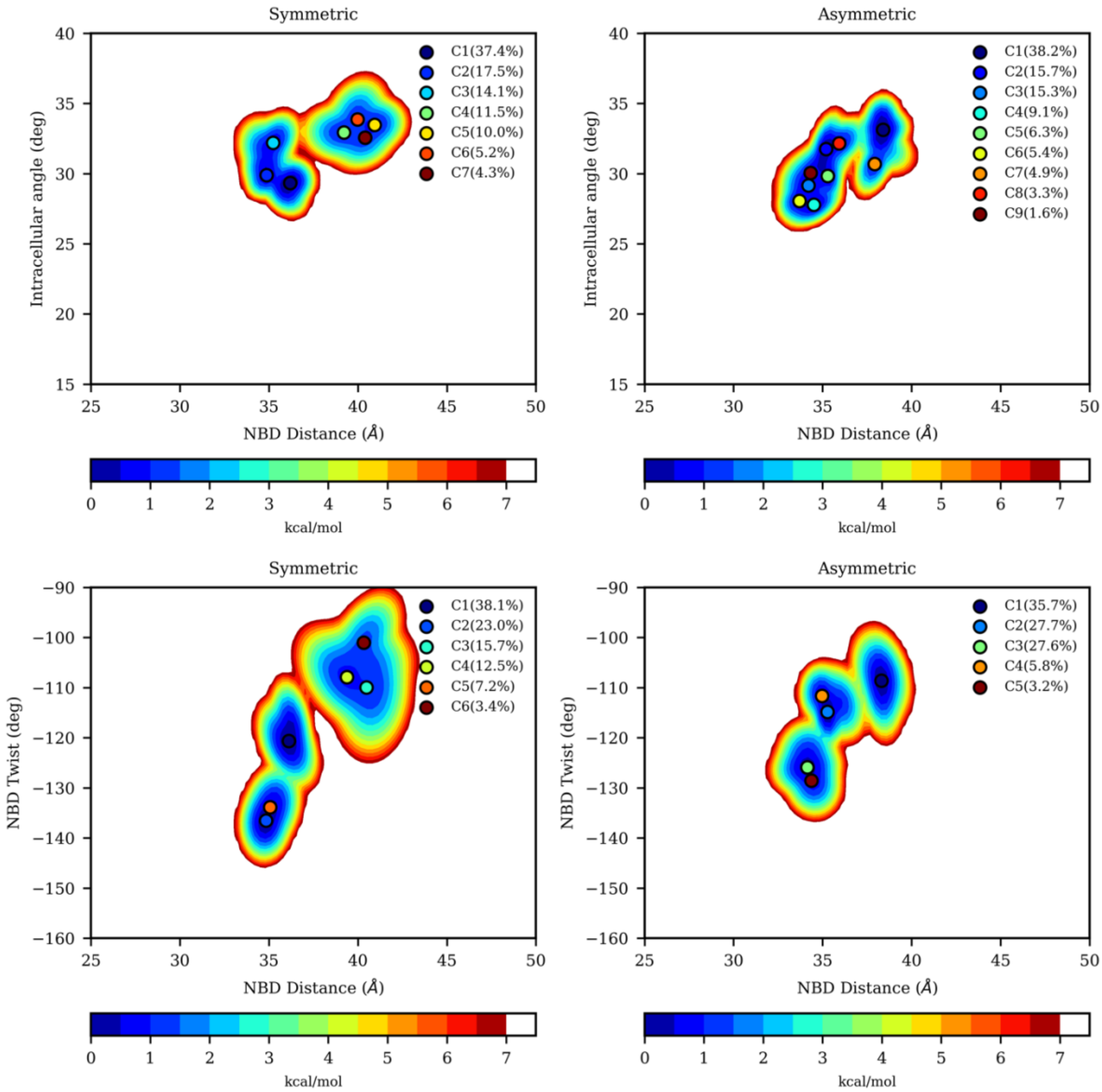

**Supporting Figure S9. Local free energy landscapes calculated for $^{IF}$ABCB4-*PC-(ATP)₂* according IC angle vs NBD distance (top) or NBD twist vs NBD distance (bottom) in symmetric (left) and asymmetric (right) membrane model from MD simulations.**

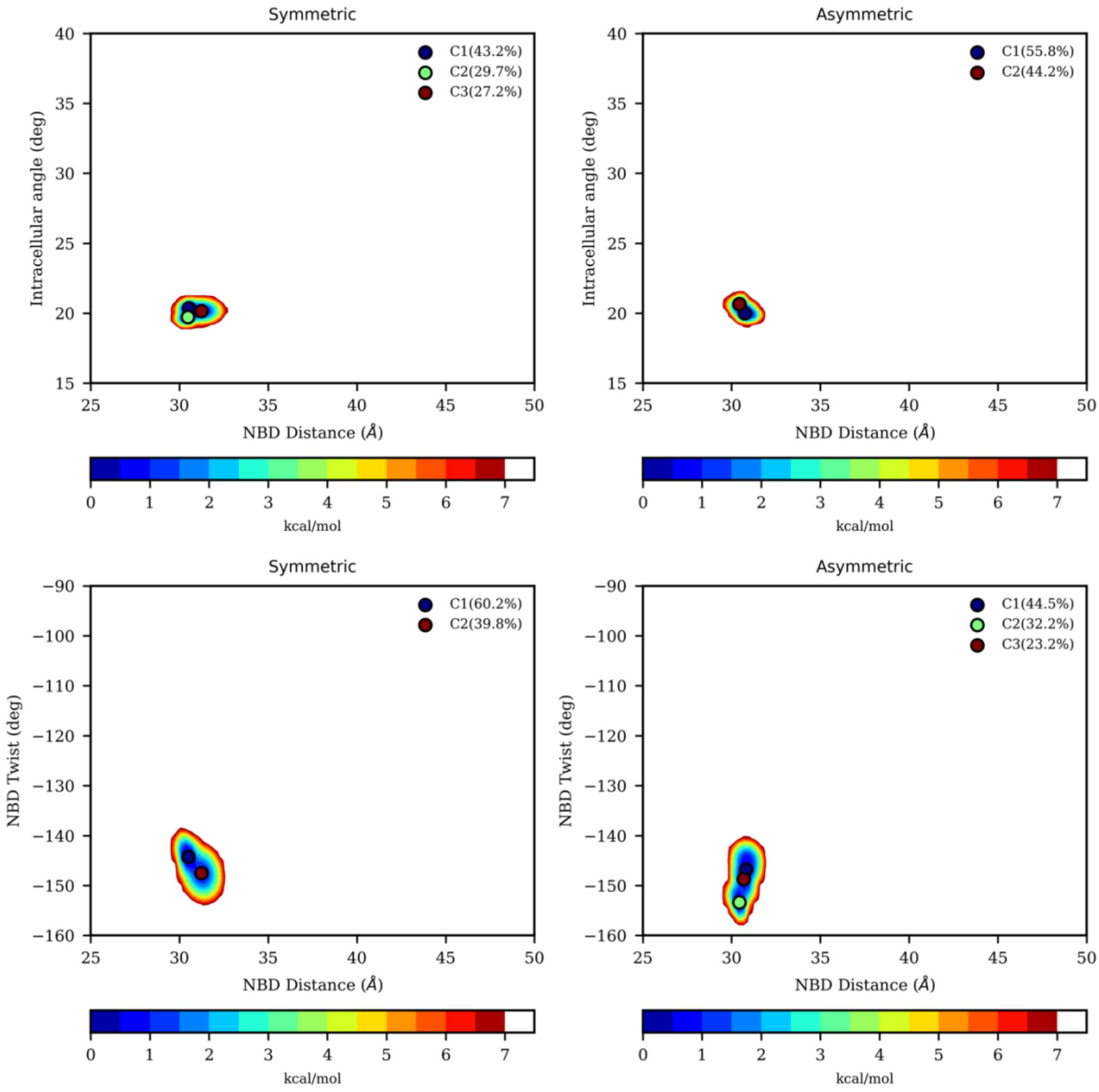

**Supporting Figure S10. Local free energy landscapes calculated for $^{CC}$ABCB4-*(ATP)*$_2$ according IC angle vs NBD distance (top) or NBD twist vs NBD distance (bottom) in symmetric (left) and asymmetric (right) membrane model from MD simulations.**

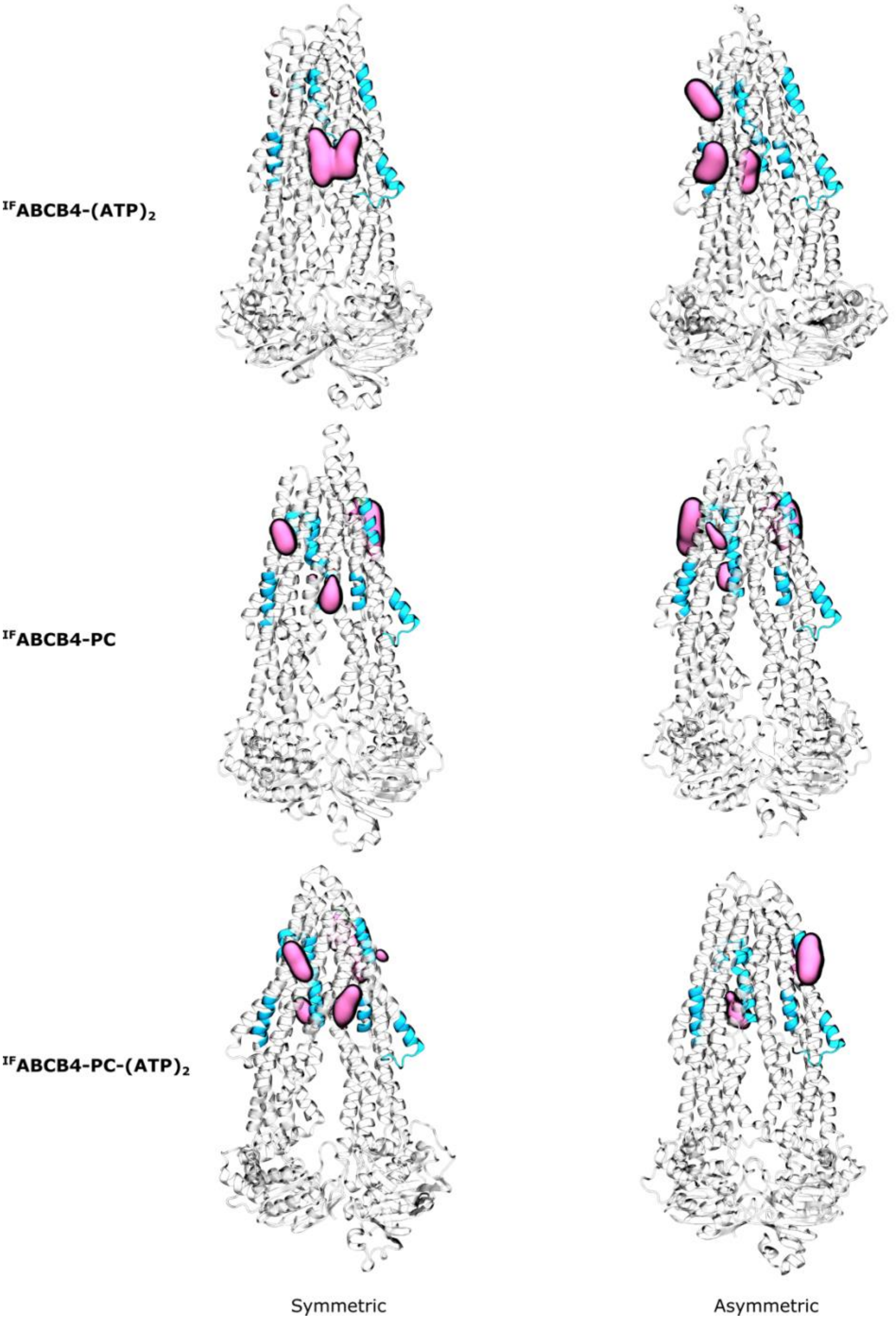

**Supporting Figure S11. Predicted cholesterol hotspots for $^{IF}$ABCB4-(ATP)$_2$, $^{IF}$ABCB4-PC and $^{IF}$ABCB4-PC-(ATP)$_2$ in symmetric and asymmetric membrane.**

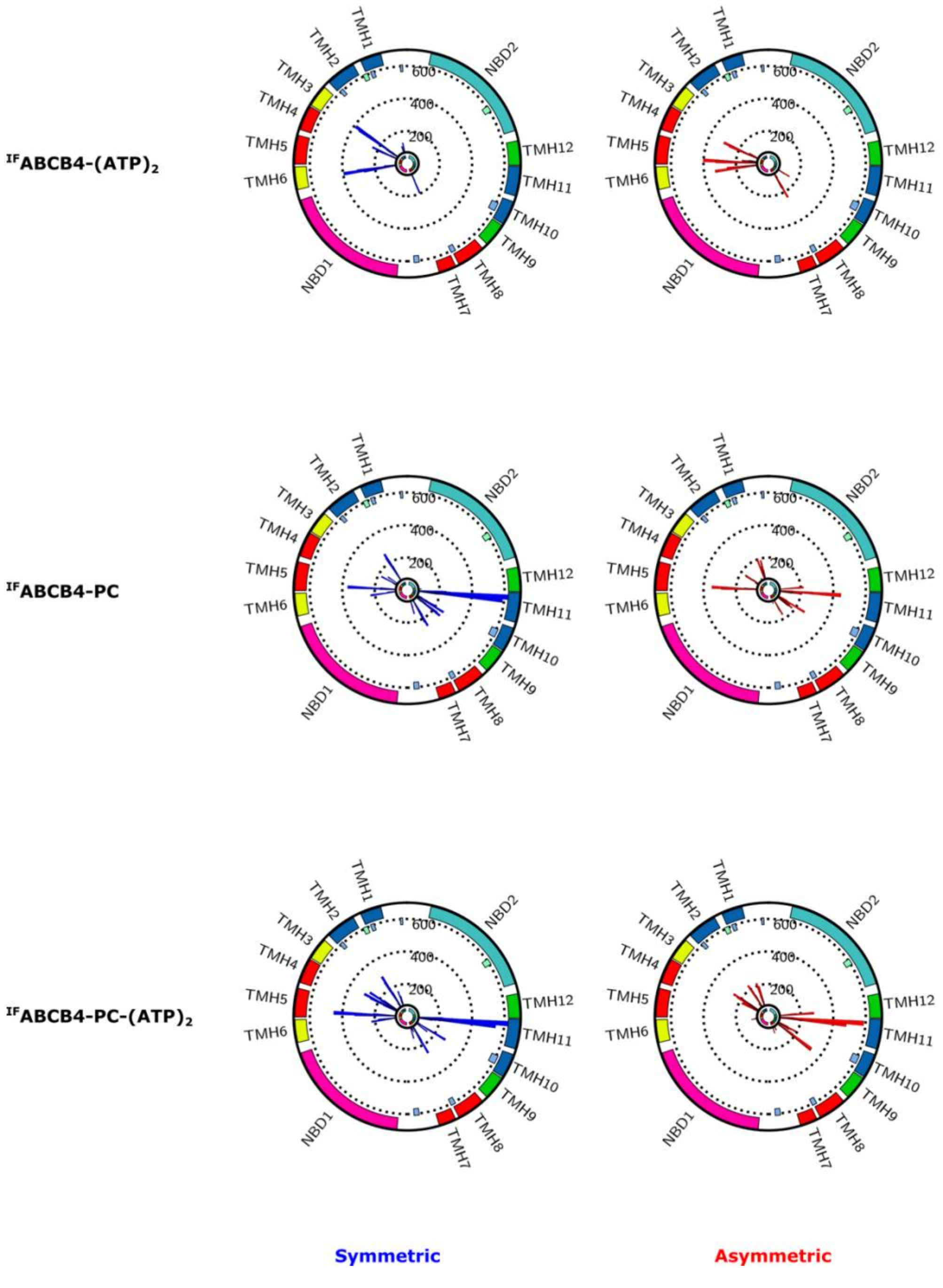

**Supporting Figure S12. Per-residue atomic contact fractions between cholesterol and $^{IF}$ABCB4-(ATP)$_2$, $^{IF}$ABCB4-PC, and $^{IF}$ABCB4-PC-(ATP)$_2$ in symmetric and asymmetric membrane models.**

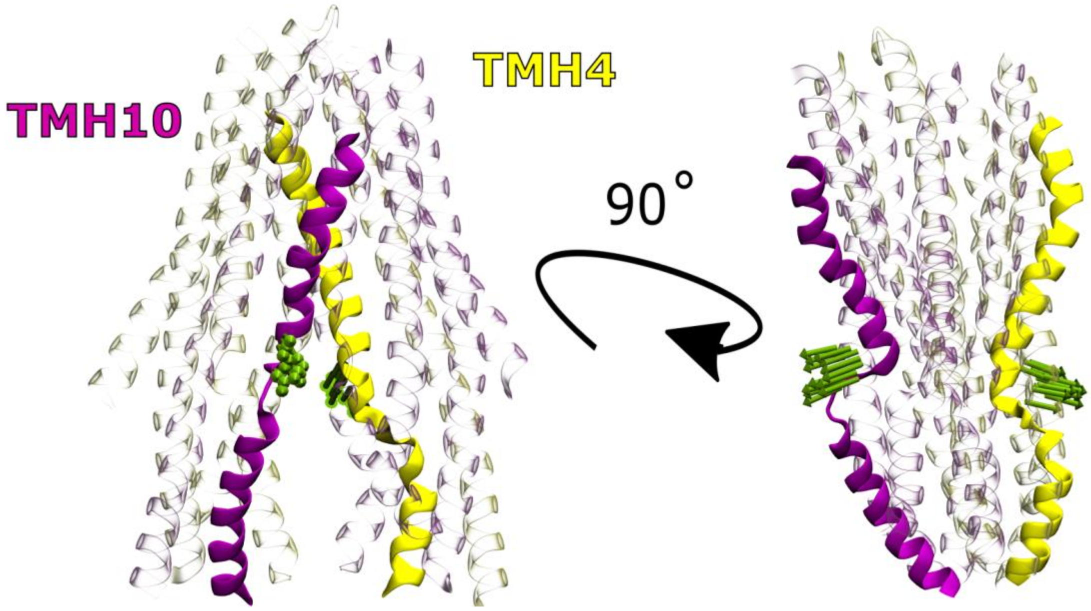

**Supporting Figure S13. Porcupine plot from PCA analyses considering all transmembrane domains of each conformation and bound state of ABCB4 in the present study highlighting TMH4 and TMH10 as main source of structural variability.**

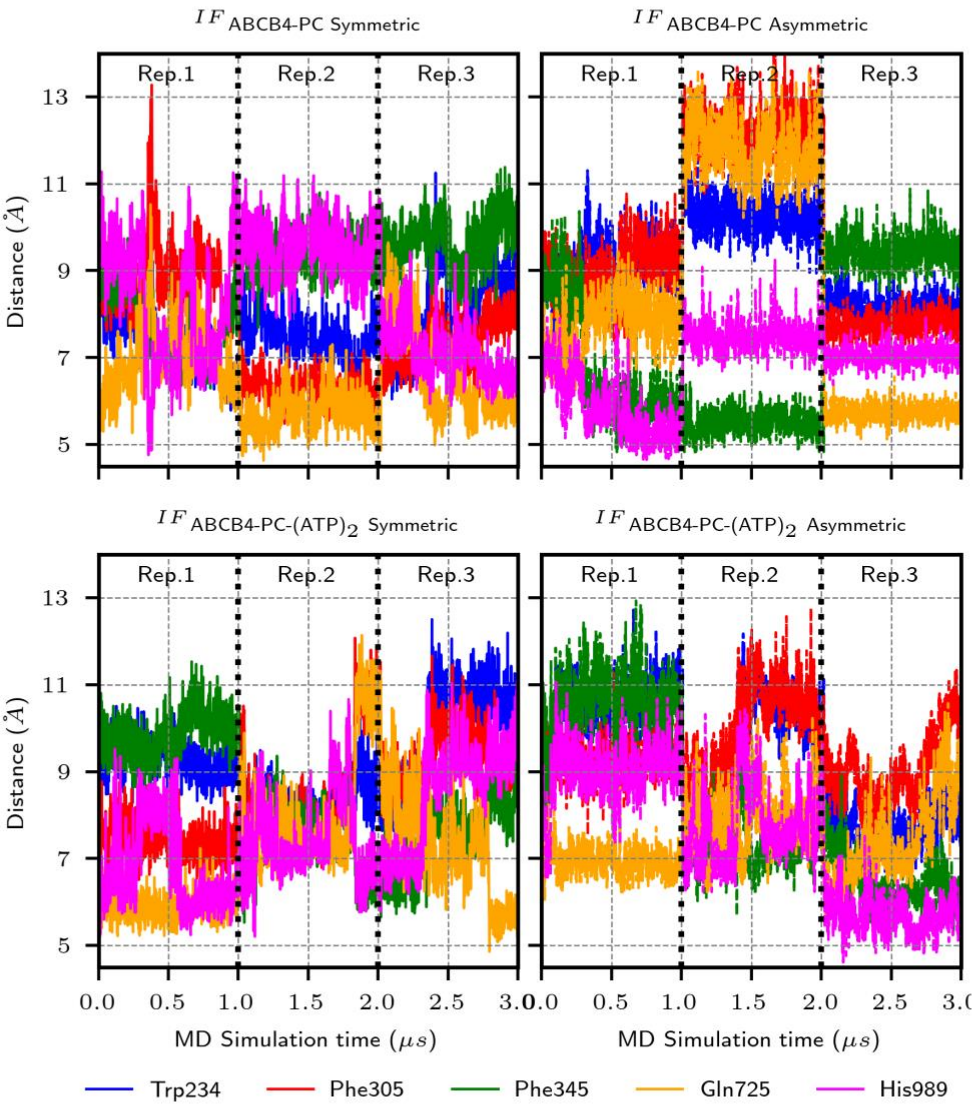

**Supporting Figure S14. Distances between substrate PC polar head and key residue identified in ABCB4 substrate binding pocket**
